# Molecular and structural remodeling of stress granules in slowly and rapidly progressive Alzheimer’s disease

**DOI:** 10.1101/2025.07.20.665556

**Authors:** Tayyaba Saleem, Matthias Schmitz, Saima Zafar, Susana da Silva Correia, Leticia Camila Flores Fernandez, Anna-Lisa Fischer, Carolina Thomas, Stefan Goebel, Wiebke Möbius, Abrar Younas, Peter Hermann, Christine Stadelmann, Olivier Andreoletti, Isidre Ferrer, Neelam Younas, Inga Zerr

**Affiliations:** Department of Neurology, University Medical Center Göttingen, Georg-August-Universitäts, Robert-Koch-Straße 40, 37075, Göttingen, Germany; German Center for Neurodegenerative Diseases (DZNE), Robert-Koch-Straße 40, 37075, Göttingen, Germany; Department of Neurogenetics, Electron Microscopy City Campus, Max Planck Institute for Multidisciplinary Sciences, Göttingen, Germany; Department of Neuropathology, University Medical Center Göttingen, Germany; Biomedical Engineering and Sciences Department, School of Mechanical and Manufacturing Engineering (SMME), National University of Sciences and Technology (NUST), Islamabad, Pakistan; UMR INRA ENVT 1225-Interactions Hôte Agent Pathogène–École Nationale Vétérinaire de Toulouse, Toulouse, France; Emeritus Professor, University of Barcelona, Barcelona, Spain; Paul-Flechsig-Institute, Center of Neuropathology and Brain Research, Leipzig, Germany

## Abstract

Stress granules (SGs) are dynamic ribonucleoprotein condensates that modulate RNA metabolism during cellular stress. Although SG dysfunction has been increasingly linked to neurodegenerative diseases, their structural and molecular remodeling in Alzheimer’s disease (AD), particularly rapidly progressive AD (rpAD), remains poorly understood. Here, we present a comprehensive multi-omics characterization of SGs from postmortem frontal cortex tissues of control, slowly progressive AD (spAD), and rpAD subjects. SGs were immunoprecipitated using Anti-TIAR antibodies and analyzed via transmission electron microscopy (TEM), LCMS/MS-based proteomics, and RNA sequencing. Key protein findings were validated in human cortical brain homogenates and a 3xTg mouse model of Aβ and tau pathology.

TEM revealed disease-specific SG morphologies: small spherical granules in controls; moderate clustering in spAD; and large, amorphous aggregates in rpAD. Proteomic profiling identified 1,667 high-confidence SG-associated proteins, including RNA-binding proteins and disease-linked proteins such as MAPT, APP, and SNCA. SGs in rpAD were significantly enriched for pathways involved in MAPK signaling, proteostasis, and neuroinflammation, while showing reduced abundance of key cytoskeletal and translational regulators, such as TUBA1B and EEF1A2.

Transcriptome analysis revealed widespread depletion of long, GC-rich, protein coding RNAs in rpAD SGs. Notably dynamic dysregulation of TUBA1B was also observed in the 3xTg mouse model and human cortical tissues, highlighting cytoskeletal vulnerability during disease progression.

Together, these findings uncover profound structural and molecular remodeling of SGs in AD, with rpAD exhibiting a distinctive shift towards pathological SG composition and function. Our results highlight a link between SG alterations and aggressive AD subtypes, providing new mechanistic insights and suggesting new potential targets for therapeutic intervention.

## 1. Introduction

Ribonucleoprotein (RNP) granules are a diverse class of membraneless organelles that assemble through multivalent interactions involving RNA-RNA, RNA-Protein, and protein-protein binding between RNPs ^1^. The discovery of mRNA-Binding proteins (RBPs) and RNA granules, along with their pivotal role in regulating the fate and function of mRNA transcripts, has highlighted the significance of translational control ^2,3^. The interaction between RBPs and RNA granules governs mRNA stability and translational activity, playing a key role in modulating protein expression both under normal and during stress conditions ^45^. The two cytoplasmic RNP granules that react to various environmental stresses are SGs and processing bodies (PBs) ^6^.

The formation and composition of SGs is specific to cell type, stress stimulus and signaling pathways. Various stresses can trigger SG formation. These include oxidative stress, heat shock, hypoxia, endoplasmic reticulum stress, nutrient deprivation, viral infection, depletion of translation initiation factors, and overexpression of RBPs ^7^. They contain 40S ribosomal subunits, translation initiation factors (i.e., eIF2, eIF3, eIF4A, eIF4G), and specific marker RBPs such as T-cell restricted intracellular antigen 1 (TIA-1), TIA-1 related protein (TIA-R), and RasGAP SH3-domain binding protein 1 (G3BP1) and 2 (G3BP2) ^8^.

SGs play a crucial role in stress response, suppressing translation and sorting mRNA for re-initiation, storage, or degradation. They may also be essential for optimizing the translation of stress-responsive and anti-apoptotic mRNAs ^9,10^. Nucleating RBPs like TIA-1, TTP and G3BP, which contain prion-like and glycine rich domains initiate SGs’ formation ^11^. As SGs mature, they incorporate microRNAs, translation initiation factors, and regulatory proteins, including kinases and GTPases. The cellular response to stress depends on its type, intensity, and duration, and it is tightly regulated by RBPs, which control global translation ^12,13^. In response to stress, SGs form in cytoplasm through liquid-liquid phase separation (LLPS) ^9,14–17^. SGs are typically temporary structures that protect cells including neurons from damaging stress. They do so by halting the mRNA translation and sequestering specific proteins and RNAs ^15,18,19^. Once the stress stimulus resolves, SGs disassemble, and nuclear proteins return to nucleus to resume translation. However, during chronic stress, SGs become persistent and transition into pathological, solid-like aggregates ^20,21^. These long-lasting SGs are associated with neurodegenerative diseases. They may act as nucleating centers for aggregation of pathological proteins, such as phosphorylated tau in neurons^22,23^. The aggregation of these RBPs resembles that of proteins associated with neurodgenerative diseases ^11^.

Neurodegenerative conditions and aging are marked by persistent oxidative stress. This activates cellular defense mechanisms that can either promote survival or trigger apoptosis. Alzheimer’s disease (AD) is a neurodegenerative disorder characterized by cognitive decline and neuron loss resulting in memory deficits, which in turn impair daily life activities. The major two hallmarks of AD are abnormal deposition of amyloid β (Aβ) plaques and neurofibrillary tangles (NFTs) of hyperphosphorylated tau ^24–26^. However, increasing evidence shows a significant variability in clinical phenotype and progression rate, raising the possibility of heterogeneous mechanisms in pathology ^27–30^. Among the pathological factors contributing to this variability, tau pathology plays a central role. The heterogeneity observed in AD may be partially explained by differences in tau propagation, Post-translational modification (PTMs) and its prion like seeding behavior, which influences disease progression and severity. Rapidly progressive AD has been recognized as a distinct clinical subtype of AD characterized by rapid progression in cognitive decline, shorter disease duration and in some cases early focal neurological signs^31,32^. While rpAD and typical AD share core neuropathological features, emerging evidence suggests that rpAD exhibit unique molecular signatures including distinct amyloid beta aggregate structures and proteomic differences in amyloid plaques.

Multiple studies suggest that environmental and physiological stress can accelerate AD pathogenesis ^33–35^. The accumulation of TIA-1 has already been reported in the brains of AD patients and in mouse model of AD ^36,37^.

Alongside disease-associated protein aggregates, numerous RNA also accumulate in SGs. This suggests that pathological SGs in AD brains may irreversibly sequester RNAs, thereby disrupting neuronal metabolism and contributing to cell loss and cognitive decline ^14,22^. However, the mechanism underlying the role of these granules in AD disease pathology and their components is unclear. In this study, we sought to investigate potential differences in the morphology and molecular composition of SGs isolated from healthy control (cont.), slowly progressive Alzheimer’s disease (spAD) and rapidly progressive AD (rpAD) human brain tissues. By analyzing the proteome and transcriptome of these granules across the disease groups, we aim to gain insights into their role in disease progression and identify any distinct alterations in granule composition or structure associated with rpAD.

## 2. Methods

### 2.1 Ethics Statement

Frontal cortex samples from 5 spAD 5 rpAD, and 5 cont. subjects were sourced from the Institute of Neuropathology Brain Bank (HUB-ICO-IDIBELL Biobank) and the Biobank of Hospital Clinic-IDIBAPS, Spain. All procedures compiled with local legislation (Ley de la Investigacion Biomedica 2013 and Real DecretoBiobancos 2014) and received approval from local ethics committees. Additionally, samples from healthy cont. subjects were obtained through the Department of Neuropathology at the University Medical Center Goettingen, Germany. Ethics approvals were granted by the Ethical Committees of the University Medical Center Göttingen, with all procedures adhering to ethical guidelines (Nr. 1/11/93 and Nr. 9/6/08)

### 2.2 Patient cohorts and spAD/rpAD Subtype characterization

The processing of patient cohorts, neuropathological assessments, and brain tissue collection for this research followed previously established protocols ^38^. Both spAD and rpAD samples exhibited Alzheimer’s disease pathologies with Braak stage ≥ V. The AD subtype samples were included without any co-existing pathologies. Detailed information on patient cohorts is provided in the supplementary Table S1. Confirmation of all the cases was done through neuropathological examination of 25 brain regions, including cerebral cortex, thalamus, diencephalon, cerebellum, and brainstem, as previously described ^39^. Various staining techniques, such as hematoxylin and eosin, Klüver-Barrera, and immunohistochemistry for glial fibrillary acidic protein, beta amyloid, phosphorylated tau, alpha synuclein, TDP43, ubiquitin, p62, and microglial-specific markers, were employed. All rpAD cases met the contemporary rpAD criteria ^31^. The Braak staging for neurofibrillary tangles (NFT) and plaque pathology, ranging from stages I-VI/0-C, as well as Thal phases (1-5).

### 2.3 Humanized 3xTg mouse model of combined A**β** and tau pathology

Oddo et al originally developed the triple-transgenic (3xTg) mouse model, which harbors mutations in APP, PS1 and tau, ^40^. In this study, 3-4 month-old 3xTg mice were stereotactically inoculated in the thalamus with 20ul of 10% (w/v) cortical brain homogenate derived from an AD patient. The homogenate was prepared in PBS and cleared by centrifugation. Both inoculated (AD) and non-inoculated (cont.) mice were sacrificed at 3, 6, 9, and 12 months decapitated and cortical tissues were rapidly collected and snap frozen in liquid nitrogen for subsequent analyses. This study was exploratory and not powered for statistical significance at each time point. A sample size of n = 4 was selected based on logistical feasibility and prior studies using 3xTg mice.

### 2.4 Isolation of endogenous stress granules

Our aim was to study the human brain-derived SGs. To achieve this aim we employed a multistep biochemical procedure, which involved three major steps: 1) Production of soluble brain lysate (S20 lysate); 2) SGs enrichment via density gradient centrifugation; (3) SGs isolation via immunoprecipitation through a monospecific affinity purified antibody. In general, keeping in view the complexity of human brain lysates, isolation of SGs from crude lysate is not possible ^41^. This multistep process is crucial to get rid of cell debris, intact cells and membrane bound organelles.

#### 2.4.1 Optiprep Density Gradient Centrifugation

To make the isolation of granules successful from human brain lysates, OptiPrep^TM^ gradient was used for density gradient centrifugation so that granule-containing fractions can be identified and selected for immunoprecipitation (*Figure **1***). Soluble S20 lysate was layered on top of 2 ml 15%-30% of OptiPrep^TM^ gradient. For fractionation of one sample, four tubes with 2 ml capacity each were used in a swinging bucket TLS55 rotor (BECKMAN 346134). After pouring the S20 lysate on top of the gradient, the lysate was subjected to ultracentrifugation at 40000rpm at 4C for 2.5 hours. After the centrifugation 12 fractions from each tube were collected from top (fraction 1) to bottom (fraction 12) carefully, and snap frozen.

**Figure 1.**
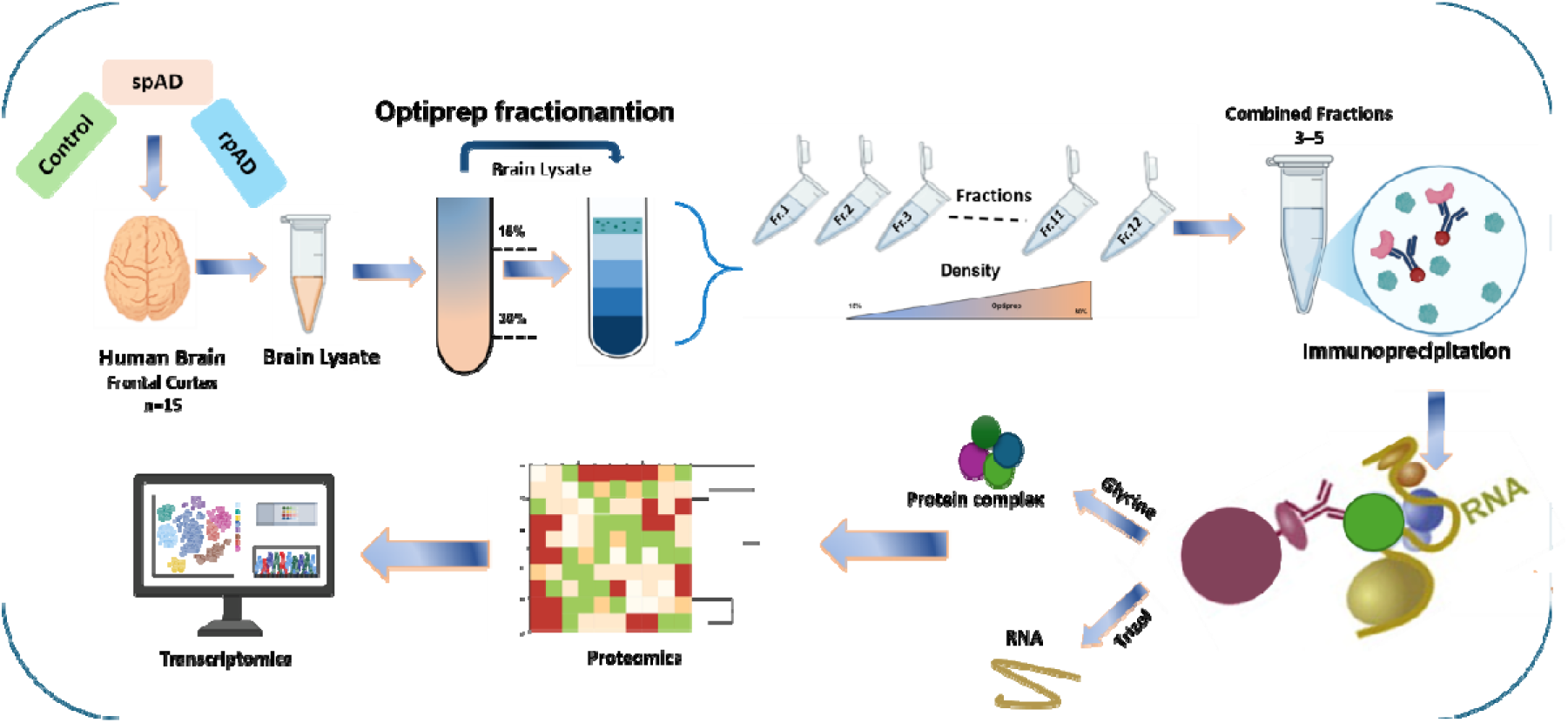
Workflow for the isolation and characterization of SGs from human brain tissue. Brain samples from cont., spAD, and rpAD cases were homogenized and centrifuged to obtain the S20 supernatant. This lysate was subjected to density gradient centrifugation using a 15–30% OptiPrep^TM^ gradient. Fractions enriched in SG markers, particularly TIA-1/TIAR, were identified and pooled. The SG-containing fractions were then subjected to immunoprecipitation using a highly specific antibody. The isolated SGs underwent further analyses, including proteomics, transcriptomics, and functional annotation to identify alterations in composition and function across disease states.

#### 2.4.2 Immunoprecipitations

After identifying the granule-enriched fractions, which were 3, 4 and 5, we proceeded for immunoprecipitation (IP). For the IP we used 200 ug of TIA-R antibody keeping in mind that it is a classical marker for SGS, and equivalent amount of Isotype control IgG was used as negative control. The antibodies were cross-linked to protein Sepharose A beads following a multistep procedure as reported before ^42^. Non-specific binding to the matrix was blocked by treating the beads with tRNA and BSA. Briefly, non-denaturing glycine elution was performed to elute the SG complexes. At the same step before elution, we separated the beads; for SGs and for transcripts associated with these granule proteins. The protein A Sepharose beads were eluted by reversible protein denaturation with glycine. The eluate was processed for mass spectrometry to find out the composition of SGs. The other half of the SG bound Sepharose beads were processed for RNA sequencing following RNA isolation.

### 2.5 Transmission electron microscopy

For TEM analysis, a 10µL sample was placed on a strip of paraffin film. A TEM grid was then carefully positioned onto the drop to allow adsorption. To enhance structural preservation, 10ul of 0.25% glutaraldehyde was added to the drop, followed by a 1-minute incubation at room temperature. The grid was subsequently washed three times in distilled water, with each wash lasting only a few seconds to remove excess fixative. Staining was performed by incubating the grid for 30 seconds in a 2% uranyl acetate solution. Following staining, the grid was air-dried without additional washing before being subjected to imaging.

### 2.6 Label-free quantification mass spectrometry (LFQ-MS) analysis

Label free quantification mass spectrometry (LFQ-MS) analysis was conducted following previously established methods ^38^. In summary, isolated SG complexes from 15 cases were separated on 4-20% Bis-tris gels (NuPAGE Novex Bis-Tris mini gels, Invitrogen) for approximately 1cm and stained using Coomassie blue. The gel bands were then cut into small cubes (1–2 mm^2^). Protein digestion into peptides, as well as their identification and quantification, followed established protocol^38^.Scaffold software (version Scaffold_4.8.4, Proteome Software Inc., Portland/OR, USA) was utilized to confirm MS/MS-based peptide and protein identifications. Peptide identification was set to a confidence level greater than 95% and protein identification required a minimum of 2 peptides with a confidence threshold of 99%. The protein prophet algorithm was used to assign protein probabilities ^43^. All the biological replicates included three technical replicates. Proteins were filtered using a fold change ≥ 1.5 compared to IgG negative controls, a threshold chosen to minimize background and enrich for high-confidence interactors.

### 2.7 RNA sequencing

For RNA isolation, we used miRNA kit and Trizol isolation. Prior to RNA extraction, all surfaces and lab equipment were treated with RNAse Zap (Thermo fisher Scientific) to prevent contamination. Total RNA was extracted from the Sepharose A beads which were already bound to the SGs and RNA by using the Trizol method. RNA concentrations were determined, and quality of the isolated RNA was assessed using a Bioanalyzer (Agilent Technologies, Santa Clara, CA).

Briefly, total RNA quality was assessed by measuring the RNA Integrity Number (RIN) using the Fragment Analyzer HS Total RNA Kit (DNF-472-FR; Agilent Technologies). RNA-seq libraries were prepared using the STAR Hamilton NGS automation platform with the Illumina Stranded Total RNA Prep, Ligation with Ribo-Zero Plus kit (Cat. No. 20040529), starting from 100 ng of total RNA. Unique dual indexes were added using the Illumina RNA UD Indexes Set A, Ligation (Cat. No. 20091655). The average fragment size of the final cDNA libraries was ∼340 bp, determined with the SS NGS Fragment 1-6000 bp Kit on a Fragment Analyzer. Library quantification was performed using the DeNovix DS-Series System. Sequencing was conducted on the Illumina NovaSeq 6000 using an S2 flow cell (100 cycles), generating approximately 25 million reads per sample.

Raw sequencing images were converted to BCL files using the Illumina BaseCaller software and demultiplexed into FASTQ files using bcl2fastq v2.20.0.422. Sequencing quality was assessed with FastQC v0.11.5.

### 2.8 Statistical analysis

All the data in this study was obtained from at least three independent experiments. Statistical analysis for mass spectrometry and RNA-Seq data was performed using Python (3.11.9), R software (R version 4.3.3), and GraphPad Prism (version 9). Libraries used in the R included DESeq2, ggplot2, pheatmap while Python included pandas, matplotlib and seaborn. Heatmap for proteomic data was created using online source Heatmapper. For comparisons between two groups, Student’s t-test or Welch’s t-tests were applied. For comparisons involving three or more groups, one way ANOVA followed by Tukey’s post-hoc test or Mann-Whitney U test was used according to normality of data. To account for multiple comparisons, we applied False Discovery Rate (FDR) correction (Benjamini-Hochberg) to control risk of type I error. Statistical significance was defined as p-value < 0.05.

### 2.9 Characterization of SGs in Alzheimer’s disease subtypes

To provide a structured overview of our approach, the schematic summarizes the key analyses conducted on SGs in cont., spAD and rpAD cases (Figure 2).

**Figure 2:**
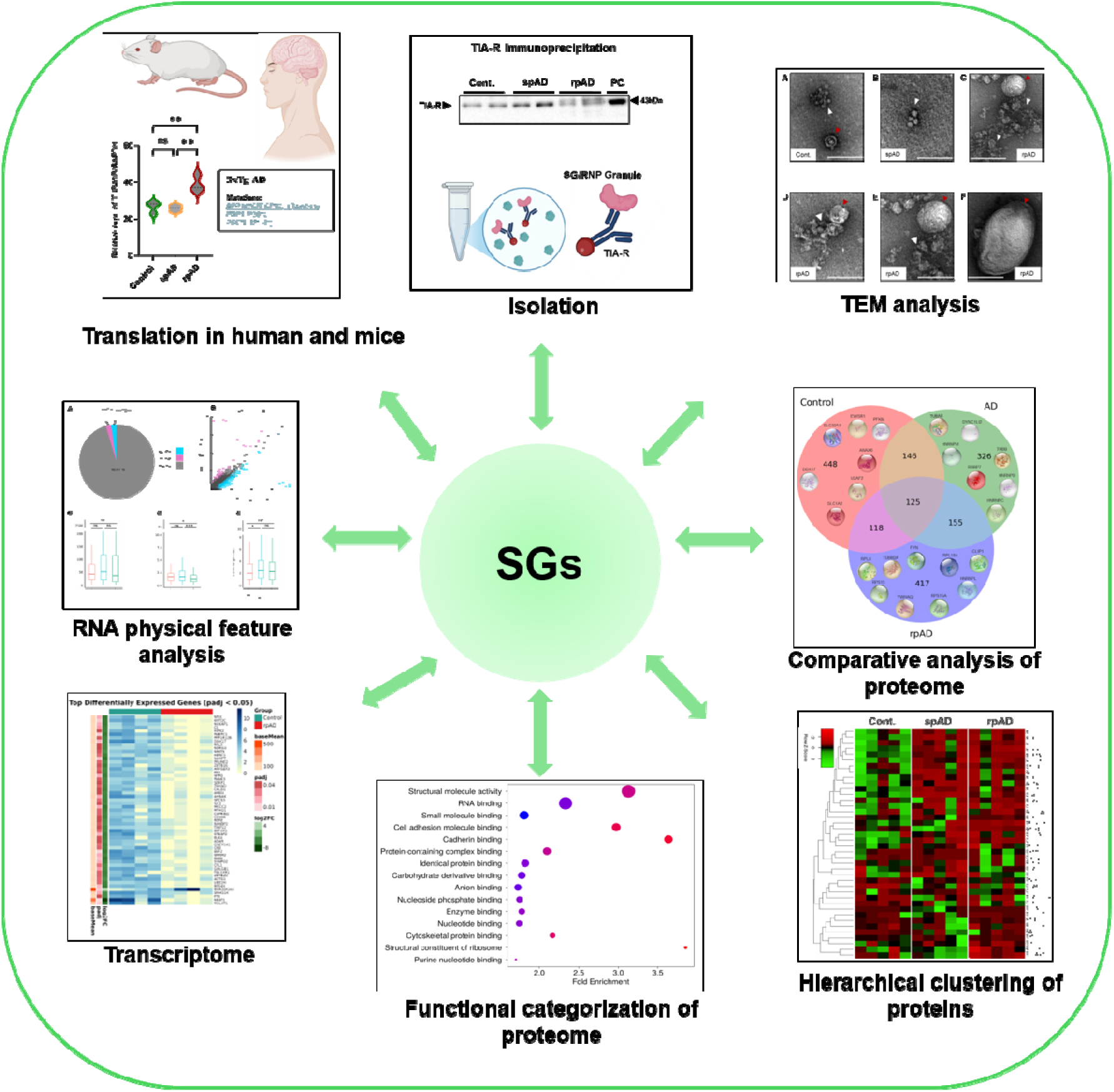
Comprehensive multi-omics characterization of SGs in cont., spAD, and rpAD. The central green node represents SGs, with surrounding panels illustrating the diverse experimental approaches used for their analysis. SGs were isolated from postmortem brain tissue using a multistep immunoprecipitation strategy targeting TIAR protein. TEM was employed to visualize SG ultrastructure and assess morphological differences among conditions. Proteomic profiling via mass spectrometry was used to identify SG-associated proteins, while RNA sequencing enabled transcriptomic analysis of SG-enriched fractions. Comparative analyses including differential expression, functional enrichment, and intersectional Venn diagram analysis were performed to uncover condition-specific alterations in SG composition and molecular function across cont., spAD, and rpAD groups.

## 3. Results

### 3.1 Characterization of SGs via immunoprecipitation using canonical markers

To isolate TIA-1/TIA-R stress granules we used a multistep protocol (Figure ***1***) ^42^. To assess the distribution of SG markers across different fractions we first analyzed the fractions using SDS-PAGE followed by Coomassie staining. The successful fractionation was confirmed by a distinct protein band pattern Figure 3 A. Western blot of the fraction was also performed with GAPDH antibody to confirm the distribution of the total protein across the fractions.

**Figure 3.**
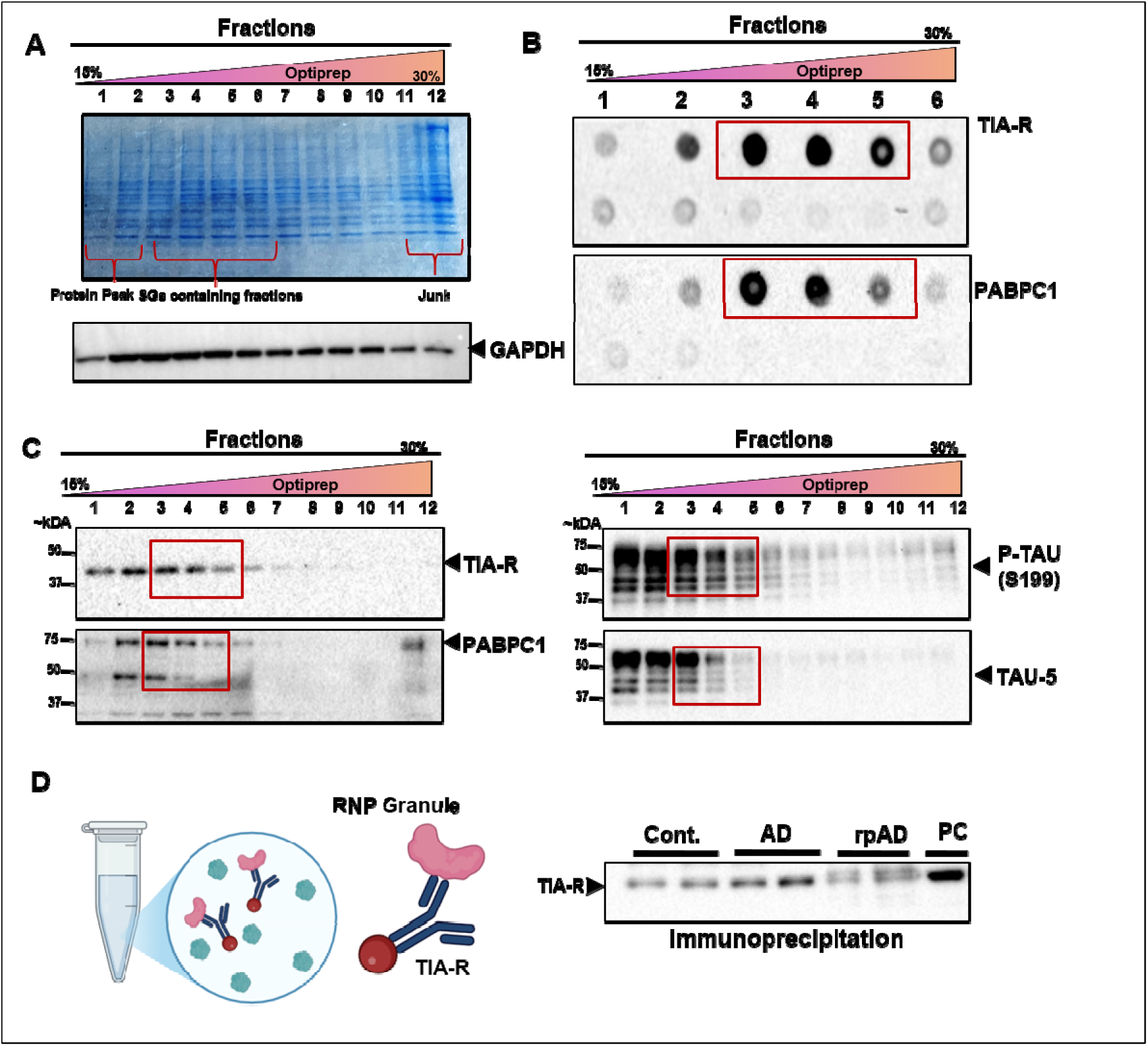
Biochemical fractionation and identification of SG-enriched fractions from human brain tissue. (A) Coomassie-stained gel of gradient fractions following density-based separation using a 15–30% OptiPrep^TM^ gradient. The first two fractions correspond to the soluble protein peak, while the bottom fractions represent debris or aggregated proteins. Fractions 3-5 are enriched in SGs, as indicated by protein distribution and subsequent analyses. GAPDH immunoblot confirms even protein distribution across fractions. (B) Dot blot analysis of non-denatured gradient fractions probed for SG marker proteins TIA-R and PABPC1 to identify native SG-enriched fractions. (C) Western blot analysis of fractions probed for TIA-R, PABPC1, phosphorylated tau (p-Tau S199), and total tau (Tau-5), confirming SG enrichment and potential tau co-association. (D) Immunoprecipitation of SG fractions using anti-TIA-R antibody confirms successful enrichment of SGs from cont., spAD, and rpAD brain samples. PC: positive control. (n= 15 (5 Cont., 5 spAD, 5 rpAD).

Next, to identify granule markers (TIA-R and PABPC1); we checked the fractions through dot blot (Figure 3B). The results showed the enrichment of granule markers in 3, 4, and 5 fractions for both TIA-R and PABPC1. The presence of TIA-R and PABPC1 in these fractions showed the possible presence of SGs, as these markers are known to associate with functional granules. The enrichment pattern observed in the dot blot provided a strong basis for selecting these fractions for downstream analysis. Additionally, the non-denaturing conditions used in the dot blot ensured the native distribution of these markers without protein denaturation artifacts.

After identifying the fractions of interest in their non-denatured state, we characterized the fractions by western blot. Western blot showed the enrichment of granule markers again in fractions 3, 4, and 5 (Figure 3C). The co-fractionation of known markers TIA-R and PABPC1 confirmed the enrichment of SGs in these fractions. Furthermore, the presence of PABPC1 suggested that these SGs were still intact and contain 3-polyadenylated mRNA. To further characterize the fractions, we examined additional SG markers and proteins implicated in neurodegeneration, such as tau-5 and p-tau S199 (Supplementary Figure S1). To investigate granule composition, fractions 3, 4, and 5 were selected for immunoprecipitation using TIA-R antibody. Keeping in view the protein peak, we excluded first two fractions even when they showed signal for the markers of interest. Given that TIA-1 and TIA-R serve as key nucleating proteins in SGs formation, this approach enabled the selective enrichment of SGs. Following multistep immunoprecipitation, the SG-enriched eluates were successfully isolated, as shown in Figure 3 D. This approach allowed us to perform downstream analyses for the investigation of their molecular composition and functional attributes. Details of dot blot analyses for PABPC1 and TIA-R across cont., spAD, and rpAD samples, along with Western blot results for FXR1, SFPQ, and oligomeric tau, are presented in Supplementary Figure S2.

### 3.2 TEM revealed increased SGs aggregation and density in rpAD

To examine the morphology of brain isolated SGs across Cont., spAD, and rpAD, we performed TEM on the isolated granules (Figure *4*). TEM analysis revealed striking differences in SG structure and organization between disease groups. According to negative staining TEM analysis, isolated SGs appeared heterogeneous, with both discrete, round-shaped structures and larger tightly packed aggregates. An overview of the SGs and zoomed-in images is presented in Figure 4. In cont. samples SGs appeared well-defined and intact predominantly maintaining spherical or near-spherical structures. No signs of adverse aggregation of SGs were observed in cont. samples. In the spAD samples, the SGs appeared smaller and discrete, potentially indicating a reduced granule formation or enhanced clearance mechanism. In contrast, rpAD samples exhibited markedly different phenotypes, with large, amorphous, and dense SGs aggregates that could indicate coalesced granules or misfolded SGs aggregates. The density of SGs and potential aggregation was high compared to cont. and spAD samples. The darker regions suggest higher electron density, which may be linked to increased aggregation of RNA-protein complexes or additional pathological inclusions.

**Figure 4:**
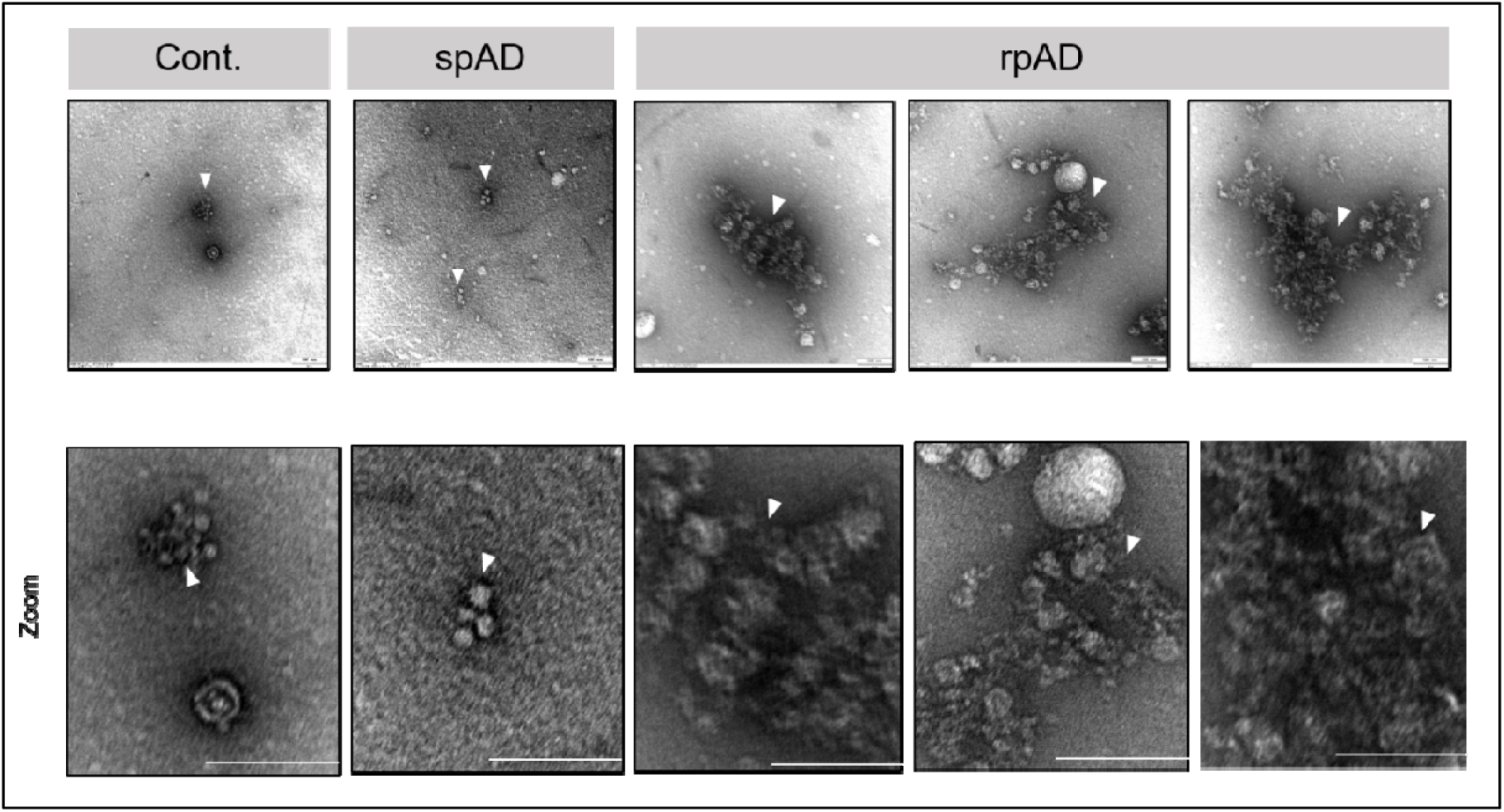
Ultrastructural analysis of SGs isolated from frontal cortex tissue of control, spAD, and rpAD cases by negative-stain TEM. Representative TEM images display SGs purified from brain homogenates of cont., spAD, and rpAD subjects. In cont. samples, SGs appear as tightly clustered, compact spherical particles (arrowheads), suggesting preserved granule integrity and organization. In spAD, SGs are more sparsely distributed, smaller, and less compact, indicative of early structural remodeling. SGs in rpAD samples exhibit highly irregular, amorphous aggregates, with disrupted morphology and apparent fusion of granules. These observations point to progressive structural disintegration of SGs across disease stages. White arrowheads indicate individual SGs or SG clusters. Scale bar: 100 nm.

The Zoomed-in images from cont. SGs showed that some granules are apparently connected through thin filamentous or string-like structures, which showed a spatial organization. In comparison to cont., SGs in spAD appeared in direct contact with each other or clustered without visible filamentous connections, which may suggest early signs of coalescence in AD SGs. Similarly, in rpAD the granules are more fused, aggregated but lacking the typical spherical structures represented by spAD SGs. These distinct morphological differences may present altered SG dynamics and clearance in rpAD.

### 3.3 Proteomic characterization of SGs in human frontal cortex

Although several approaches have been employed to study the SGs in the disease state and in cell models, the molecular composition of these granules from AD is not yet understood ^44–46^. To characterize the proteomic composition of isolated SGs we performed LC-MS/MS analysis from frontal cortical region of cont. spAD and rpAD subjects. Each group consisted of five biological replicates, with three technical replicates per sample. In total, we identified 3250 proteins in our SG dataset. To map the SGs composition, we first compared the identified human brain-isolated SG proteins from our MS dataset with curated SG protein databases known as RNA granule database and Mammalian Stress Granule Proteome (MSGP) database. Of the 3250, proteins identified in our study 566 overlapped with a curated RNA granule database (1354 proteins) which contains reported proteins of SGs from human cell lines. An overlap of proteins from our study with the RNA granule database is presented in Supplementary Figure S3. Additionally, when matched to the MSGP database that contains 464 proteins, our dataset showed an overlap of 266 proteins. The significant overlap suggested that our data captured a good portion of known SG-associated proteins while potentially identifying novel candidates as well. To refine the datasets and enhance the specificity we applied stringent criteria of filtering the dataset with negative control IgG data. The proteins for SGs proteome analysis were retained only if the fold change was ≥ 1.5, compared to IgG negative control. After filtering the background, noise and potential contaminants, 1,667 proteins were retained across all groups (cont. spAD, and rpAD) ensuring the high confidence proteins enrichment for downstream analysis.

### 3.4 Hierarchical clustering revealed distinct SG granule proteomic Profile in AD subtypes

SGs from human brain form a dense network of protein-protein interactions. Among the 417 uniquely enriched proteins in rpAD group, HNRNPUL, RBBP9, HNRNPL, EEF1B2, SERPINH1, RPS13, RPS12 were more prominent based on their role in SG biology (Figure 5 A) Through high throughput technique mass spectrometry, we identified several proteins, which are crucial components of SGs such as CAPRIN1, PABPC1, TIA-1, G3BP1, DDX6, HNRNP, EIF4A3. Among 448 uniquely enriched proteins in cont. group were PRR4, ANXA6, DDX17, and PLA2G2A etc. Moreover, the 326 uniquely enriched proteins in the spAD group represented SERPING1, FAM162A, APOL1, HNRNPAB, EIF5, PRDX3 and RBM3.

**Figure 5:**
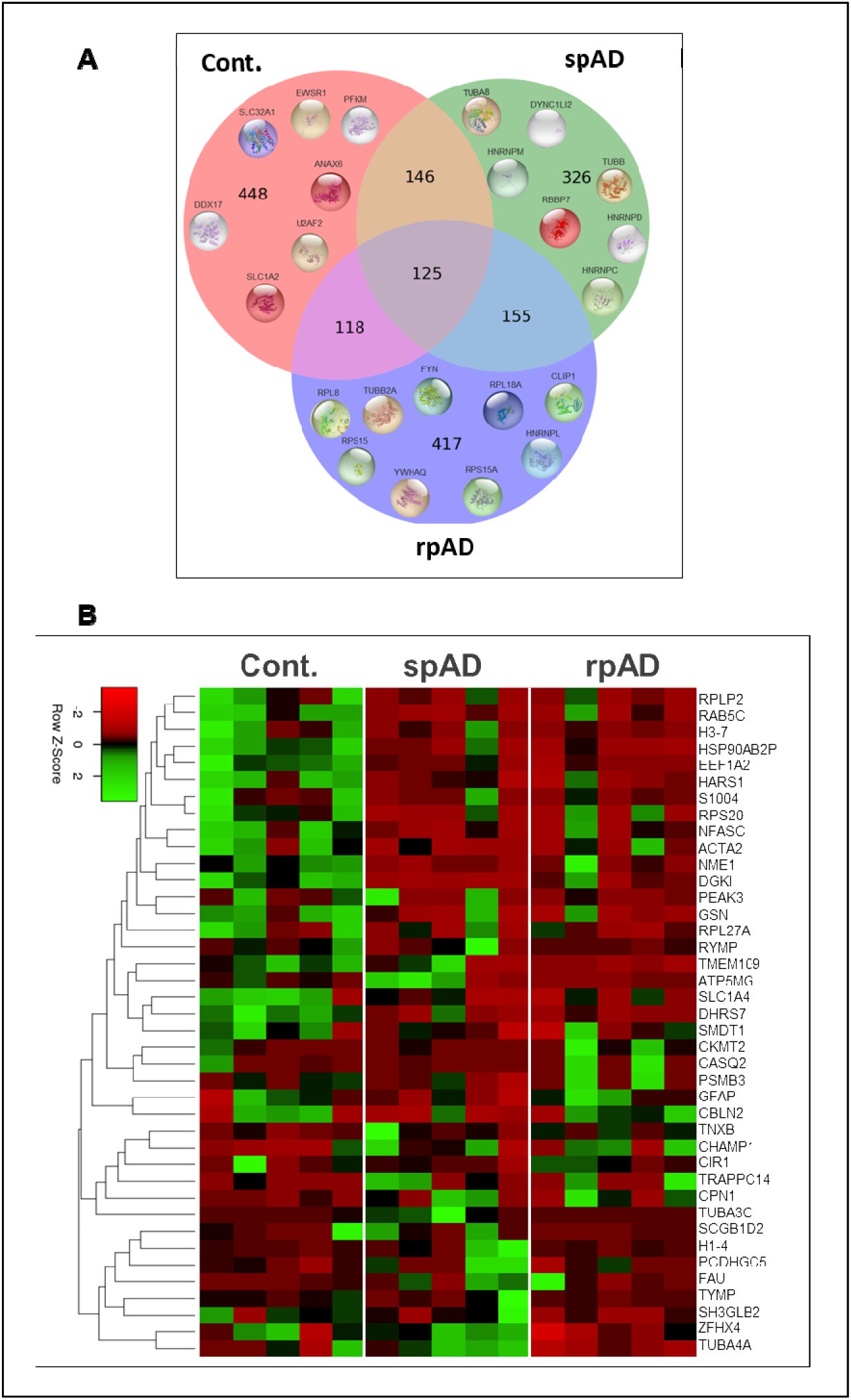
Differential SG proteome profiles across cont., spAD, and rpAD. (A) Venn diagram illustrating the overlap and uniqueness of SG-associated proteins identified by mass spectrometry in cont. (n = 5), spAD (n = 5), and rpAD (n = 5) brain samples. A total of 125 proteins were commonly detected across all groups. Cont. samples contained 448 unique proteins, while spAD and rpAD groups showed 326 and 417 unique proteins, respectively. Selected condition-specific proteins are highlighted with corresponding structure images. (B) Heatmap of differentially enriched proteins (DEPs) from SG proteomic analysis. Log□-transformed protein intensities were Z-score normalized across all samples. Each column represents an individual biological replicate, and each row corresponds to a significantly altered SG-associated protein. Green indicates higher abundance, and red indicates lower abundance. A total of five biological samples and three technical replicates were included per group.

Hierarchical clustering analysis showed distinct disease-specific proteomic patterns across cont., spAD and rpAD groups. The heatmap showed the significantly modified proteins across the groups (Figure 5B). Heat Shock Protein 90 Alpha Family Class B Member 2, Pseudogene, (HSP90AB2) is found significantly less enriched in spAD and rpAD SGs’ proteome compared to cont. group. We found significantly less enrichment of HARS1, NFASC and DGKI in the spAD and rpAD SGs compared to cont. group shown by our MS data from SGs. Majority of these proteins were part of the RNA granule database and differential abundance of these components might indicate dysregulation in SG dynamics.

Interestingly the analysis from the proteins, which were only found in spAD and rpAD SGs proteome, showed that TUBA1B was less enriched in the rpAD group compared to spAD. ZFHX4 and PLCB3 were also found less enriched particularly in the rpAD group when quantified against spAD. The proteins which were only present in the SGs proteome from cont. and rpAD group were also quantified from the MS results. RPS14, GSN, GNAS, TMEM109 and YBX1 were found significantly less enriched in the rpAD proteome as compared to cont. group. Interestingly all of these proteins are part of RNA granule database. Precise role of these proteins in the disease pathogenesis is not clear yet however the underrepresentation of these proteins in rpAD might contribute to impaired SGs function. Similarly, the comparison of the SGs proteome from cont. and spAD showed the less enrichment of HSPA1B, NME1, STX1B, SEPTIN7, FXYD6 and DPYSL2 in the spAD group (Supplementary Figure S4).

Interaction network analysis of the proteins which were differentially enriched across the disease groups from our SGs proteome showed a strong interaction network given by STRING However, ZFHX4, FXYD6, TMEM109, GNAS showed no interaction at all (Supplementary Figure S5).

### 3.5 Dysregulation of TUBA1B and EEF1A2 in human brain and 3xTg mouse model

MS analysis of SGs revealed differential abundance of cytoskeletal and translational proteins across cont., spAD, and rpAD cases. TUBA1B showed less enrichment (p=0.0079) in rpAD SGs compared to spAD (Figure 6A). EEF1A2 was also found less enriched (p=0.007) in both spAD and rpAD compared to cont. (Figure 6B). Western blot validation in human cortical brain homogenates showed the presence of three distinct isoforms or cleavage products (250 kDa, 52 kDa, and 35 kDa) with disease specific variation in band intensity. The overall expression of TUBA1B was found upregulated in rpAD compared to cont. and spAD (Figure 6A). TUBA1B was less abundant in SGs from rpAD but increased in total brain homogenate, possibly reflecting compensatory upregulation or redistribution from SGs to the cytoskeleton during stress. To extend the findings beyond terminal-stage human pathology, expression of TUBA1B was analyzed in 3xTg mice at 3, 6, 9, and 12 mpi. TUBA1B protein levels varied over time, with higher expression observed at early time points (pre-symptomatic stage) and decreased levels at later stages (symptomatic phase) (Figure 6D). EEF1A2 expression was also evaluated in human brain tissues and mouse model. No prominent difference in EEF1A2 expression were observed in the 3xTg mouse model; however, a significant downregulation (p=0.0085) was detected in rpAD human brain tissue consistent with the SG proteomic data (Figure 6E). EEF1A2 was consistently downregulated in human rpAD brain and SGs but not in 3xTg mice, suggesting possible species-specific regulation or model limitations.

**Figure 6.**
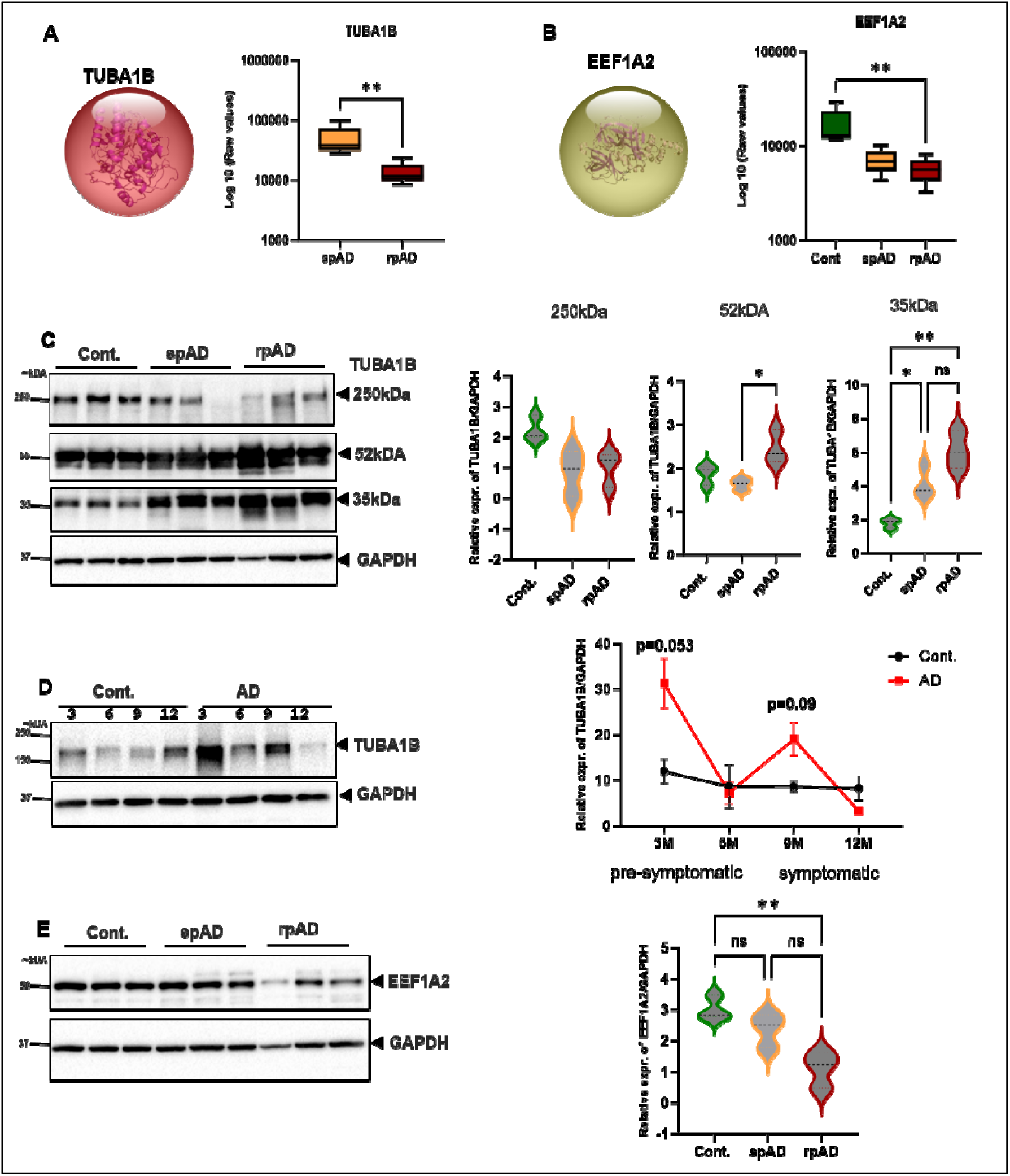
Differential expression and validation of TUBA1B and EEF1A2 in human SGs and brain tissue from AD subtypes and mouse models. (A) Mass spectrometry analysis of SG revealed increased abundance of TUBA1B in spAD and rpAD compared to cont. samples (n = 5 per group). Statistical significance assessed using the Mann-Whitney U test. (B) EEF1A2 was significantly reduced in SG fractions from rpAD compared to spAD and cont. (n = 5 per group). (C) Western blot validation of TUBA1B in human cortical brain homogenates (n = 3 per group) revealed distinct disease-associated expression profiles. Quantitative analysis confirmed significant group differences. (D) Temporal analysis of TUBA1B expression in cortical brain tissue from 3xTg-inoculated AD mice (n = 4) and age-matched non-inoculated controls (n = 4) at 3, 6, 9, and 12 months post-inoculation (mpi). Statistical comparisons at each time point were performed using unpaired t-tests with correction for multiple comparisons. (E) Western blot analysis of EEF1A2 expression in human brain tissue showed significant downregulation in rpAD (n = 3 per group). Statistical comparisons across three groups (Panels B, C, E) were performed using one-way ANOVA with Tukey’s post hoc test. *p□<□0.05; **p□<□0.01; ***p□<□0.001; ****p□<□0.0001.

### 3.6 GO enrichment analysis of SG proteome

The GO term analysis of the filtered data (SG proteome of human brain tissue from present study) showed the enrichment of neurodegenerative pathways and disorders such as Parkinson’s disease (PD), AD, Huntington’s disease, amyotrophic lateral sclerosis (ALS), and Prion disease (Figure 7A). Top biological processes enriched were related to SGs function such as cytoplasmic translation (Figure 7B). The overrepresentation of terms like secretory granule, RNA-binding proteins, stress response factors and translation regulators in the data highlights that isolated granules are enriched with SG-related core components as shown in (Figure 7C-D)

**Figure 7:**
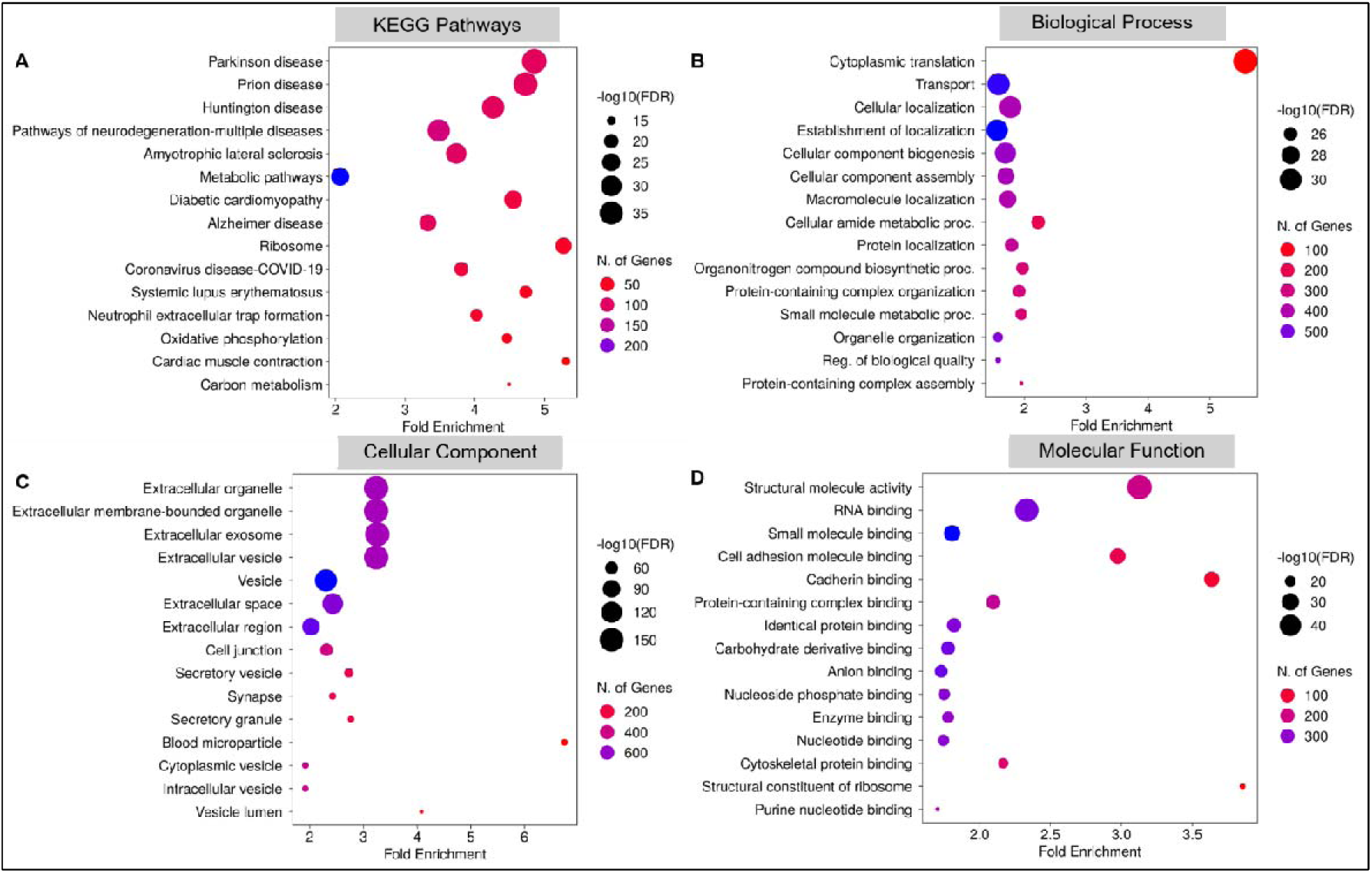
Functional enrichment analysis of stress granule (SG)-associated proteins across multiple annotation categories.(A) KEGG pathway enrichment highlights significant overrepresentation of neurodegenerative disease pathways, including PD, AD, Huntington’s disease, ALS, and Prion disease, among SG-associated proteins. (B) GO biological process terms show enrichment for pathways related to translation, localization, component assembly, and regulation of protein quality. (C) GO cellular component analysis identifies predominant localization of SG-associated proteins to extracellular organelles, exosomes, vesicles, synapses, and secretory granules. (D) GO molecular function analysis reveals enrichment for RNA binding, structural molecule activity, protein complex binding, and nucleotide-binding activities. In all panels, dot size represents the statistical significance of enrichment as −log□□ (FDR), and color intensity indicates the number of overlapping genes. FDR values were calculated using the Benjamini-Hochberg method. Fold enrichment indicates the degree to which each term is overrepresented relative to background.

#### 3.6.1 Pathological rewiring of SGs in AD enhances MAPK and inflammatory signaling

Functional enrichment analysis revealed a stark contrast between the protein constituent uniquely detected in cont. SGs versus those found exclusively in spAD and rpAD. In cont. SGs, predominantly enriched biological processes were involved in ion homoeostasis, DNA repair, ribosomal function, and translational control. These functions align with the canonical role of SGs in maintaining neuronal proteostasis and facilitating recovery from cellular stress by transiently repressing translation and sequestering damaged mRNAs or proteins ^47,48^. Conversely, SGs isolated from spAD and rpAD brains exhibited enrichment for proteins linked to MAPK signaling, lipoprotein metabolism, and acute inflammatory responses, suggesting a shift in SG identity under neurodegenerative conditions (Figure 8). Notably, components associated with lipoprotein particles and immune activation such as apolipoproteins and complement related proteins were detected supporting the hypothesis that SGs may become entangled in maladaptive inflammatory signaling cascades during disease progression ^49,50^. This functional divergence supports a model in which SGs transition from a localized, neuroprotective role in healthy neurons to a more extracellular, pro-inflammatory entity in AD pathology. This remodeling could contribute to enhanced neuroinflammation, vascular dysfunction, and the propagation of tau pathology, in line with recent studies highlighting the role of SGS in tau aggregation and neurodegeneration ^51,52^. These findings emphasized that SGs in spAD and rpAD are not merely markers of cellular stress but may actively participate in disease progression through altered protein recruitment and impaired resolution dynamics.

**Figure 8.**
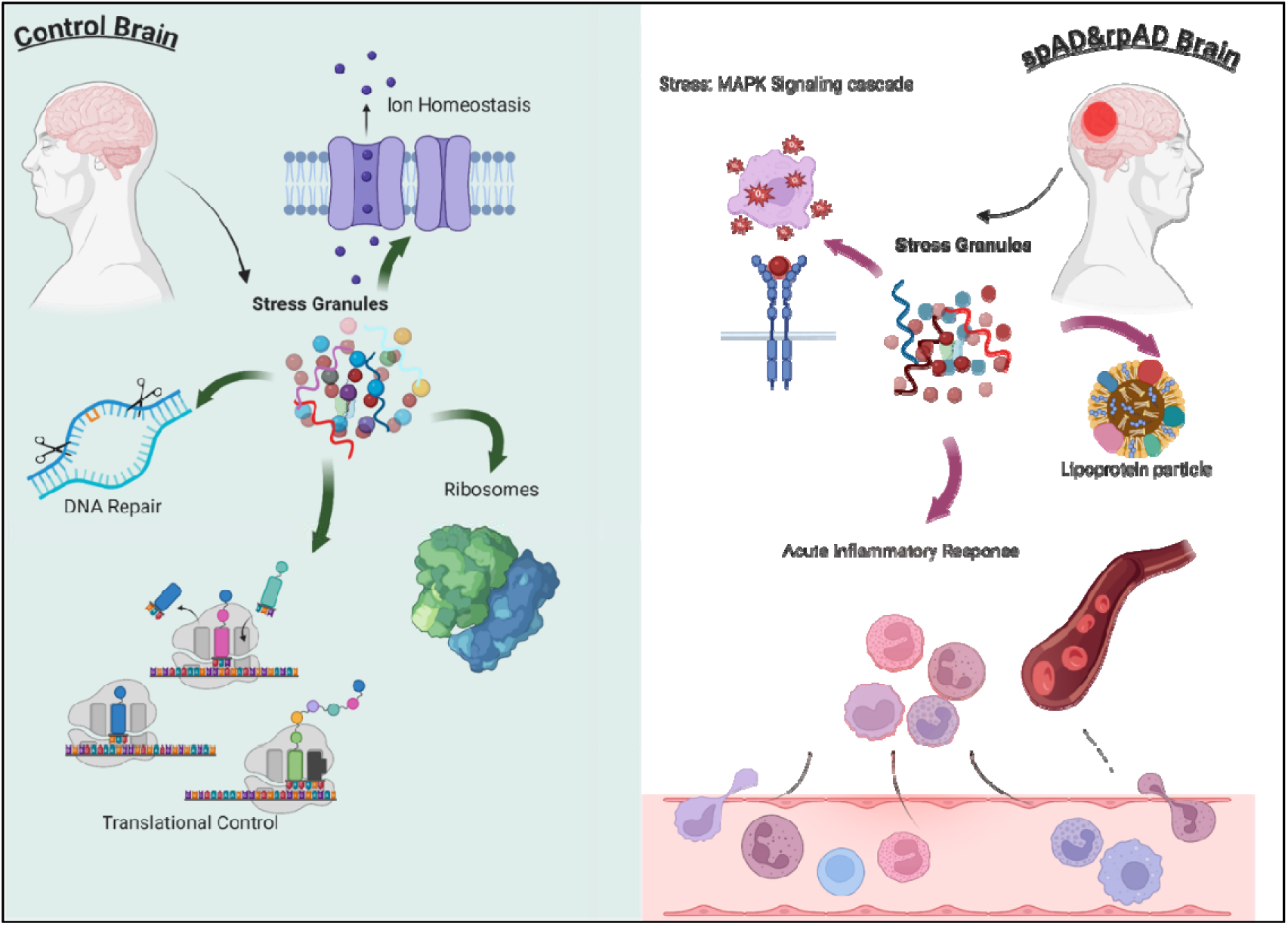
Functional divergence of SGs between healthy and diseased brains. In the cont. brain (left), SGs contribute to neuronal homeostasis by supporting ion homeostasis, DNA repair, ribosomal integrity, and translational control. These SGs remain intracellular and serve protective, regulatory roles during cellular stress. In contrast, in the spAD and rpAD brains (right), SGs exhibit pathological remodeling. Stress activates the MAPK signaling cascade, leading to the formation of aberrant SGs. These SGs are associated with lipoprotein particles and might be implicated in driving an acute inflammatory response, potentially through extracellular release or interaction with immune pathways. This functional shift marks the transition from beneficial SG regulation in healthy tissue to maladaptive, pro-inflammatory signaling in Alzheimer’s disease subtypes.

### 3.7 Transcriptome of SGs in Alzheimer’s disease subtypes

We then characterized RNA component of granules to understand complete repertoire of cont., spAD and rpAD, as these condensates contain RNA and proteins. RNA-sequencing data was acquired from RNA isolated from the immunoprecipitated complexes; RNA attached to RNA binding proteins. We identified 17541 transcripts in total including transcripts identified in single biological replicate. RNA-seq libraries from one cont. and one rpAD sample failed quality control due to RNA degradation (RIN < 6), and were excluded from further analysis. Differential analysis later excluded all the transcripts that were not detected in 60% of the samples in any of the groups.

The biotype of transcriptome of the SGs from human brain revealed the similar pattern of enrichment of different RNAs as reported from the SGs transcriptome of cells^53^. Protein coding mRNA constituted the major portion of the transcriptome followed by long non-coding RNAs (lncRNAs) as shown in Figure 9A.

**Figure 9:**
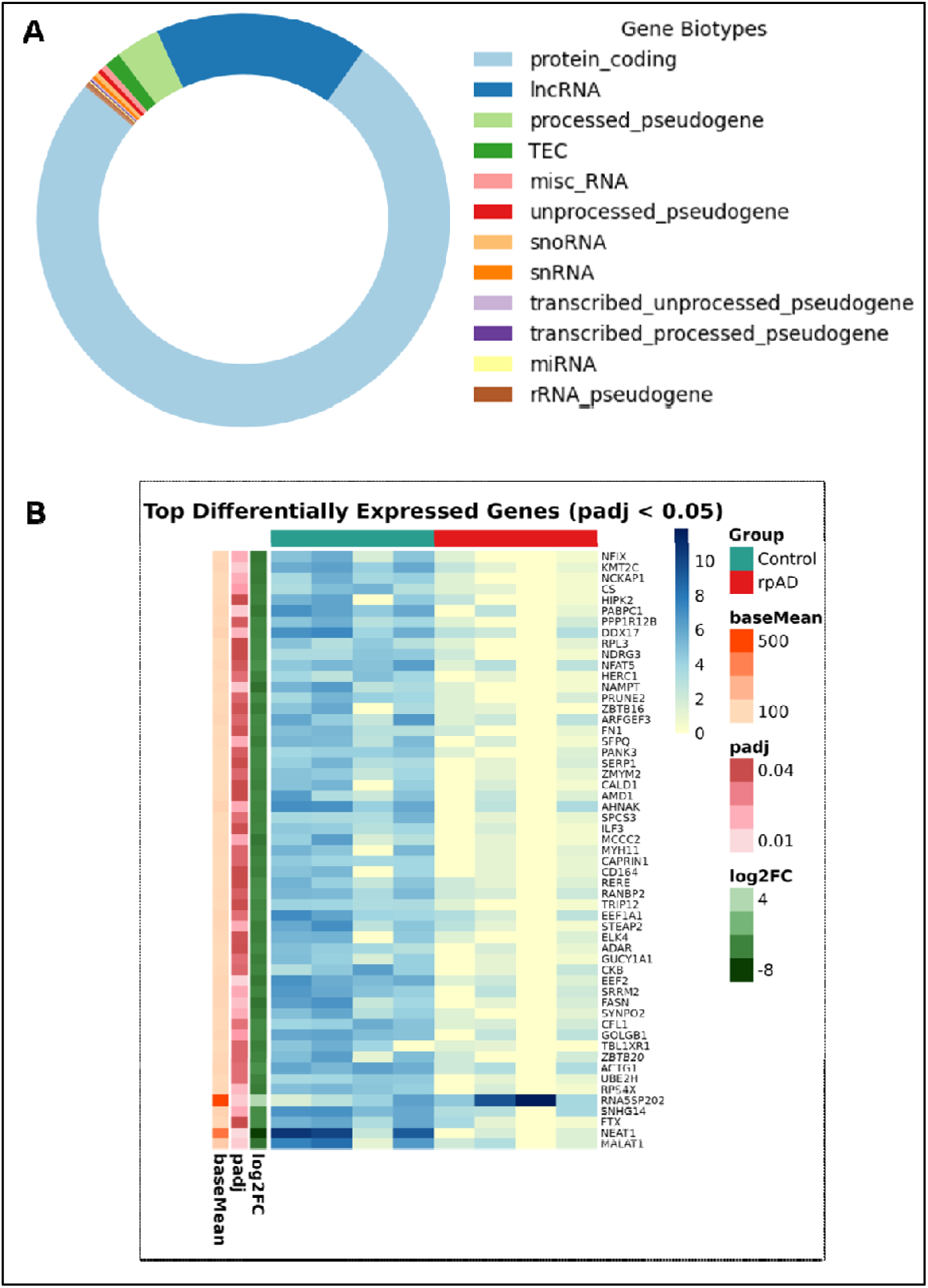
Transcriptomic analysis of stress granules (SGs) isolated from human brain tissue. (A) Donut chart showing the distribution of gene biotypes within the SG-associated transcriptome. The majority of transcripts belong to the protein-coding category, with contributions from lncRNAs, pseudogenes, miRNAs, and other RNA species such as snRNA, snoRNA etc.(B) Heatmap of the top 55 differentially expressed genes (DEGs) between cont. and rpAD groups (adjusted p-value < 0.05) based on DESeq2 analysis. Rows represent individual genes, and columns represent samples from cont. (teal) and rpAD (red) groups. Color intensity indicates normalized expression levels (variance-stabilized counts), with blue representing higher expression and yellow representing lower expression. Accompanying gene-level annotations include log□ fold change (log□FC, green scale), adjusted p-values (padj, red scale), and mean expression level (baseMean, orange scale). Sample size: n = 13 total (Cont. = 4, spAD = 5, rpAD = 4).

Differential enrichment analysis from cont. vs rpAD showed the depletion of NEAT1, CSDE1, NFIX, and enrichment of DDX5, KCNH1 and BCYRN1. Analysis of spAD vs rpAD showed the enrichment of CSDE1, VPS13C, NEAT1 etc. in the spAD SGs transcriptome.

Heatmap of top transcripts, which were differentially enriched across the groups, is also presented in Figure 9B. The analysis of rpAD and cont. showed that majority of the proteins which were less enriched in the rpAD group were RBPs and part of RNA granule database. EEF1A1, DDX17, CAPRIN1, GSK3B, MALT1, PABPC1, and SFPQ were of more importance given their role in the AD and neurodegeneration. Transcriptome data from the SGs proteome from the human brain highlighted the transcripts, which were contained in the granules for longer times and maybe disrupting the normal function of these proteins consequently disrupting major processes in the cell leading to the disease.

### 3.8 Physical properties of RNA differed among *depleted*, *enriched*, and *neither* category in disease subtypes

To explore whether physical properties of RNA are associated with their differential abundance in disease, we analyzed transcript characteristics based on their enrichment or depletion patterns. Differential expression (DE) analysis was carried out to compare the transcript levels across cont. vs. spAD, cont. vs. rpAD and spAD vs. rpAD using DEseq2. Transcripts were grouped into three categories in each of three DE analysis based on their fold-change thresholds. Specifically, transcripts were labelled as *enriched* (fold change>2), *neither* (0.5 ≤fold) and *depleted* (fold change <0.5). To assess robustness, we tested alternative cutoffs (FC > 1.5 or <0.5), which yielded similar trends in rpAD depletion, albeit with expanded transcript lists.

To explore further, we assessed the transcript features including transcript length, coding sequence length (CDS), 3’UTR length, 5’UTR length and, GC content among *enriched*, *depleted* and *neither* category. In spAD vs rpAD scatter plot, striking overrepresentation of *depleted* transcripts in rpAD (n=1582, blue dots) was observed compared to minimally *enriched* transcripts (n=10, pink dots) suggesting global transcriptional suppression in rpAD. Gray dots present *neither* category with no significant changes in expression. Moreover, we found that *depleted* transcripts were significantly longer than *neither* transcripts (p<0.001) which suggests that longer transcripts are preferentially downregulated in the rpAD. Similarly, *depleted* transcripts showed significant longer CDS length compared to *neither* (p<0.001). We further explored 5′UTR and 3′UTR length (Figure 10B).

**Figure 10:**
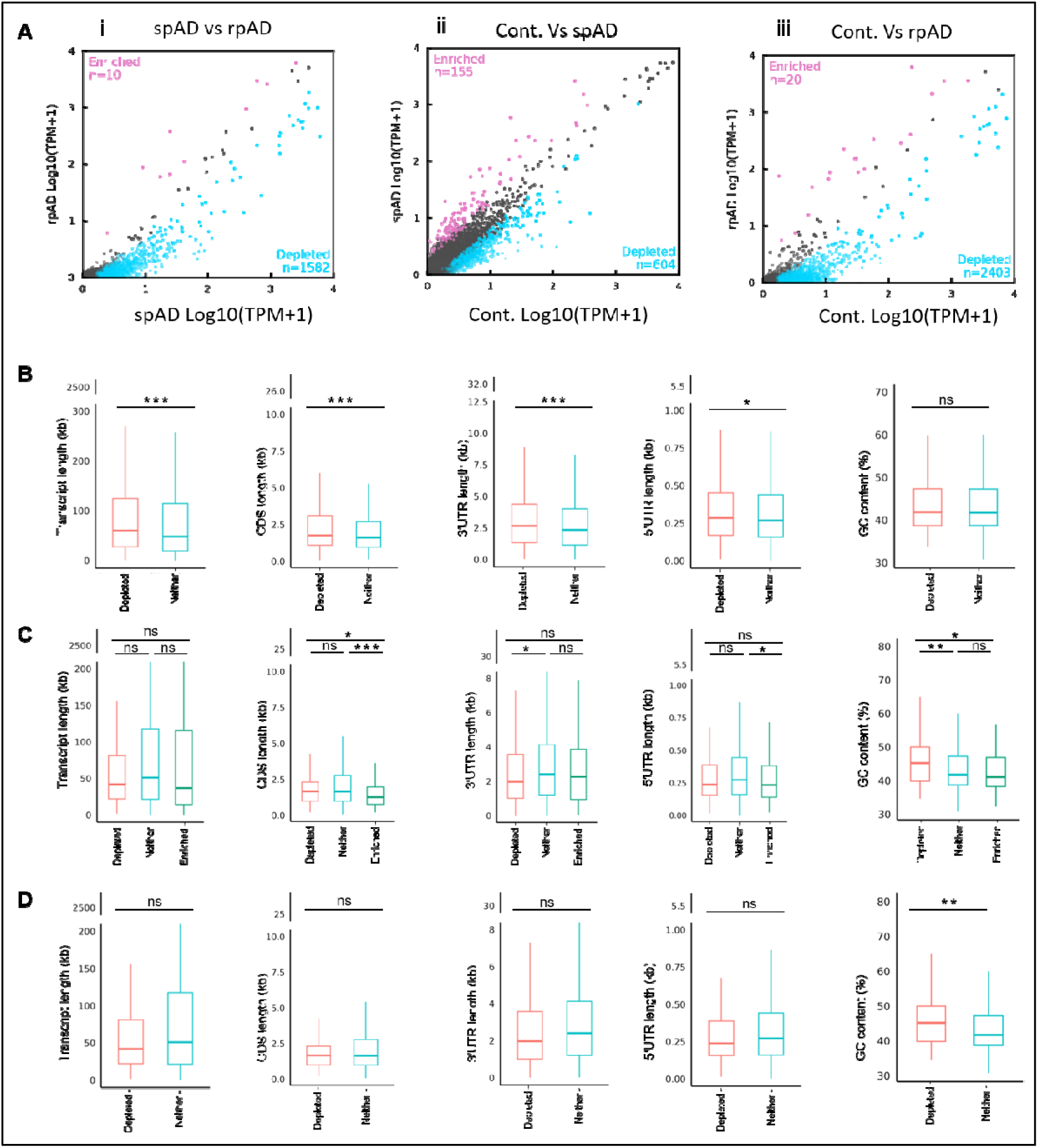
Comparative analysis of RNA abundance and transcript properties in stress granules (SGs) from spAD, rpAD, and control brain samples. (A) Scatter plots showing transcript abundance comparisons across groups based on TPM values (Log10(TPM+1)). Pink dots represent *enriched* RNAs (fold change >2, TPM >1), blue dots represent *depleted* RNAs (fold change <0.5, TPM >1), and dark gray dots represent unchanged RNAs (“*neither*”). (A (i) spAD vs rpAD (A (ii) Cont. vs spAD (A (iii) Cont. vs rpAD (B) Comparison of physical RNA properties between *enriched*, *depleted*, and *neither* categories in spAD vs rpAD. Boxplots show differences in total transcript length, coding sequence (CDS) length, 3′UTR length, 5′UTR length, and GC content. (C) Same analysis as in (B), applied to cont. vs spAD. (D) Same analysis as in (B), applied to cont. vs rpAD. Statistical significance was calculated using the Wilcoxon rank-sum test: *p < 0.05; **p < 0.01; **p < 0.001; ns = not significant. Outliers were included in the statistical test but truncated from y-axis display for better visualization. Sample size: n = 13 (Control = 4, spAD = 5, rpAD = 4).

Keeping in view that these lengths are crucial metrics for SG aggregation, we found that 3′UTR length of *depleted* transcripts was significantly longer than *neither* transcripts (p<0.001). Similarly, 5′UTR length of *depleted* transcripts was also significantly longer than in the *neither* category (p<0.05).

5′UTR length can decrease translational efficiency by reducing translation initiation because of RNA structure and preferential binding to small non-coding microRNAs ^54^. We found no significant difference for GC content between *depleted* and *neither* transcripts (*Figure 10B*). We then compared the physical features of transcripts between spAD compared to cont. The scatter plot showed that *depleted* transcripts (n=604) were more common than *enriched* transcripts (n=155) (*Figure 10B*). No significant differences were observed for the transcript length between all three categories However, CDS length varied significantly across *enriched* and other categories. *Enriched* transcripts showed significant shorter CDS length compared to *neither* and *depleted* (p<0.0001) indicating that shorter protein coding genes tend to be sequestered in spAD while slightly shorter CDS transcripts are more likely not to be captured and depleted from SGs. For UTR analysis, we observed that *enriched* transcripts have longer 3’UTR compared to *neither*, while *enriched* and *depleted* transcripts showed no significant differences in terms of 3′UTR length. The analysis of 5′UTR length showed significant difference between *enriched* and *neither* transcript with shorter 5′UTR transcripts present in *enriched* group. Lower GC content was observed in *enriched* transcripts compared to *depleted* ones (*Figure 10C*). On the other hand *depleted* transcripts had higher GC content compared to *neither* (p<0.01).

Finally to assess specific physical properties of RNA that contribute to differential transcript abundance between cont. and rpAD SG we analyzed GC content, transcript length, and coding sequence length. Scatter plot shows overrepresentation of *depleted* transcripts in rpAD (n=2403) compared to *enriched* transcripts (n=20) again suggesting global transcriptional suppression particularly in rpAD (*Figure 10A* iii.). Unlike spAD vs rpAD comparison, transcript length did not show any significant difference between *depleted* and *neither* category. Similarly, CDS length was not varied between *depleted* and *neither* transcripts in cont. vs. rpAD comparison. In the same way 5′UTR and 3′UTR length were not significantly different between *depleted* and *neither* categories. However, GC content between *depleted* and *neither* categories significantly varied (p <0.01) (*Figure 10D*). *Depleted* transcripts showed lower GC content compared to *neither*.

### 3.9 Transcriptomic profiling of SGs: Predominance of SG biology pathways

To attain a deeper understanding of SGs transcriptome at functional level we performed GO enrichment analysis. This analysis showed key biological processes, and molecular pathways associated with the SGs transcripts. Pathway analysis of transcriptome showed the enrichment of neurodegeneration related pathways exactly like proteome data that includes AD, Parkinson’s disease, and Huntington’s disease. Additionally, transcripts were enriched in dopaminergic synapse and prion disease pathway, further connecting the role of SGs to neurodegeneration. Other enriched pathways included ribosomes and protein processing in endoplasmic reticulum. For the biological process, we found the overrepresentation of protein localization and protein related metabolic pathways that aligns with the translational aspect of SGs. The enriched terms for cellular component showed cytoplasmic vesicles, focal adhesion and extracellular vesicles in our data. Moreover, analysis of molecular functions revealed strong enrichment of mRNA-binding, RNA-binding, protein specific domain binding, and protein containing complex binding, underscoring the importance of SGs’ role in RNA metabolism and post-transcriptional regulation (supplementary Figure S6). Additionally, transcription factor binding, and ubiquitin-protein ligase binding were significantly enriched highlighting the involvement of SG-associated transcripts in protein degradation under cellular stress. These are the most important aspects of SGs biology as RNA binding to the RBPs play a role in formation of the granules and actin filaments binding is a crucial part of cytoskeleton that plays a vital role in the SGs’ dynamics.

#### 3.9.1 rpAD Is characterized by distinct disruption of proteostasis, RNA processing, and cell adhesion pathways

To uncover biological processes and pathways that may differentiate between spAD and rpAD, we performed GO and KEGG pathway analysis on DEGs between the clinical subtypes of AD. Pathway analysis showed significant enrichment of cellular adhesions and immune relevant processes, with key pathways including focal adhesion, adherens junction and ubiquitin-mediated proteolysis (Figure 11A). The presence of ribosomal and spliceosome pathways showed probable altered RNA processing in rpAD. Notably, viral carcinogenesis, proteoglycans in cancer and microRNAs in cancer were also found enriched emphasizing potential dysregulation of cellular stress responses and signaling pathways that may aggravate the disease progression in rpAD. Similarly, significantly enriched biological processes included RNA splicing, negative regulation of DNA damage and cell cycle checkpoints, and metabolic processes pointing out subtype specific disruption in cellular stress adaptation and signaling pathways (Figure 11B). DEGs identified in rpAD compared to spAD were significantly associated with extracellular vesicles and synaptic structures featuring ribonucleoprotein complexes, focal adhesion sites, exosomes, and cell junctions (Figure 11C). Interestingly, molecular function analysis revealed the strong enrichment of protein binding activities, involving misfolded protein binding; ubiquitin-protein ligase binding and RNA binding, highlighting potential dysregulation in proteostasis processes that are crucial in neurodegenerative progression (Figure 11D). The enrichment of enzyme and hydrolase activity, phosphate inhibition and nucleic acid binding further suggests widespread disruption of metabolic pathways in rpAD compared to spAD. To complement our main findings, we also conducted functional enrichment analysis of differentially expressed transcripts between cont. and rpAD groups; detailed results are available in Supplementary Figure S7.

**Figure 11:**
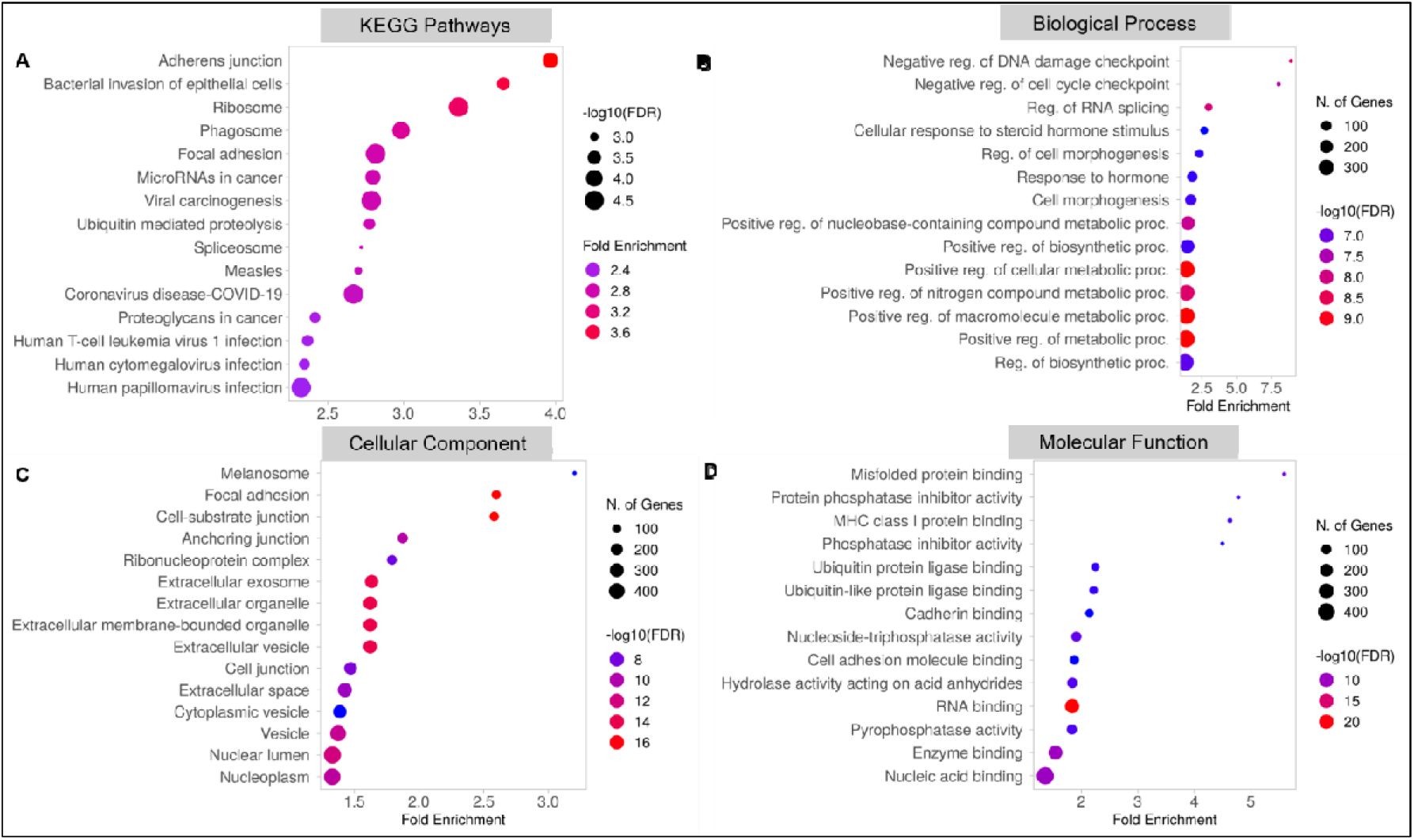
Functional enrichment analysis of DEGs in SGs between spAD and rpAD brain samples. (A) KEGG pathway enrichment analysis reveals significant overrepresentation of pathways such as adherens junctions, ribosome, phagosome, viral carcinogenesis, and spliceosome among DEGs from the SG transcriptome. (B) GO biological processes enriched among DEGs include negative regulation of DNA damage checkpoint, RNA splicing, cellular response to hormone stimuli, and regulation of biosynthetic and metabolic processes. (C) GO cellular component analysis identifies predominant transcript localization to ribonucleoprotein complexes, exosomes, extracellular vesicles, and anchoring junctions. (D) GO molecular function terms show enrichment for binding activities such as misfolded protein binding, MHC class I protein binding, RNA binding, and hydrolase and ligase activities. Dot size represents statistical significance as -log□□(FDR), while dot color indicates the number of genes associated with each term. Fold enrichment reflects the magnitude of overrepresentation compared to background. FDR values were calculated using the Benjamini-Hochberg method via ShinyGO.

## 4. Discussion

SGs are dynamic ribonucleoprotein condensates that modulate RNA translation and protein homeostasis in response to stress ^47,55^. While SG dysfunction is increasingly linked to neurodegeneration, its role in AD, and particularly its rapidly progressive form (rpAD), remains poorly characterized. Here, we provide the first integrative analysis of SGs isolated from postmortem human brains across cont., spAD, and rpAD groups, using TEM, proteomics, transcriptomics, and in vivo validation in a 3xTg AD mouse model.

### 4.1 Structural and Molecular Remodeling of SGs in rpAD

Our TEM analysis demonstrated marked structural differences in SG morphology across disease groups. Cont. SGs appeared as small, spherical condensates consistent with transient granules described under physiological stress. In contrast, rpAD SGs were larger, amorphous, and aggregated, phenotypes that mirror pathological granules seen in ALS and tauopathies ^56,57^. These morphological changes suggest impaired SG clearance, potentially due to autophagy dysfunction, which has been reported to exacerbate SG accumulation and drive tau aggregation ^51^. SG formation is typically a reversible physiological process, regulated by signaling cascades that enable assembly and disassembly in response to cellular stress. However, under prolonged stress conditions, this physiological aggregation can become pathological. It has been proposed that mutant and aggregation prone proteins interact with RBPs within SGs leading to impaired SG disassembly ^58^. This pathological shift could contribute to persistent SG accumulation in rpAD, thereby exacerbating disease progression. Our findings align with neuropathological studies in AD, which reports that large SGs are characteristic of microglial cells in the advanced disease stage, suggesting that persistent stress stimuli drive SG retention ^59^. Moreover, another study demonstrated that SGs, particularly those containing RBPs such as TIA-1 and TTP localize with tau pathology as AD progresses ^52^. To further investigate the molecular underpinnings of these morphological differences, we performed proteomic profiling of brain-derived SGs.

Proteomic profiling identified over 1,600 high-confidence SG-associated proteins across all groups. Compared to cell-line models, our dataset revealed a significantly broader repertoire, likely reflecting the complexity of brain-derived granules. Interestingly, our data revealed that SGs contain many proteins that neither possess prion-like domains nor directly bind to RNA. This aligns with prior findings indicating that, in addition to typical SG markers, other unexpected proteins are also incorporated into the SG proteome ^60^. In our opinion, these proteins could be the interaction partners of the primary SGs marker and may influence the assembly of RBPs in the granules or promote the aggregation. Traditional cell-based research often employs acute stressors, such as sodium arsenite, to induce SG formation, leading to identification of a limited set of proteins. In contrast, the persistent nature of SGs from the human brain in our study may recruit more protein diversity. Indeed, it is already reported that prolonged heat shock stress induces compositional and structural changes in SGs. Specifically, during the transition from acute to prolonged stress, SGs initially increase in size while their number decreases, followed by a stabilization phase where SGs size remains unchanged ^61^.

Known RBPs, including CAPRIN1, G3BP1, TIA-1, and DDX6, were abundant, along with cytoskeletal and chaperone proteins. Quantitative analysis revealed that the SG proteome in rpAD exhibited reduced enrichment of TUBA1B, EEF1A2, HSPA1B, and SEPTIN7, suggesting that SG composition shifts from protective to dysfunctional under chronic stress. These proteins are integral to cytoskeletal dynamics and translation control, both essential for SG resolution ^48,62^. Building on this broader proteomic landscape, we next focused on the enrichment patterns of well-characterized SG-associated proteins.

### 4.2 SG-Associated Cytoskeletal Dysfunction in rpAD

Among the proteomic alterations, cytoskeletal components emerged as critical differentiators between rpAD and cont. conditions, prompting a closer examination. We observed significantly lower enrichment of TUBA1B and TUBA4A in rpAD SGs. Both encode αtubulin isoforms critical for microtubule stability. Loss-of-function variants in TUBA4A are linked to ALS and frontotemporal dementia (FTD), and downregulation in our rpAD dataset mirrors findings in FTD brain tissue ^63^. A stable cytoskeleton is essential for SG disassembly; its disruption can lead to persistent granules and impaired cellular recovery. The biphasic expression of TUBA1B in 3xTg mice; early elevation followed by decline, further supports dynamic cytoskeletal stress during disease progression. Western blotting of human tissue validated the presence of TUBA1B with increased levels in rpAD brain homogenates possibly reflecting compensatory upregulation or aggregation. Interestingly our proteome data revealed the presence of tau and its associated isoforms in the SGs further supporting the interaction of RBPs and tau in SGs. Tau-F (2N4R) has been found significantly downregulated in the rpAD SGs compared to cont., which suggests a 3R/4R balance favoring microtubule instability.

### 4.3 Translation Machinery Is Disrupted in SGs and Brain Tissue

In addition to cytoskeletal instability, components of the translation machinery were also notably disrupted. EEF1A2, a multifunctional elongation factor with chaperone and RNA-binding capacity, was significantly downregulated in rpAD SGs and human cortex. EEF1A2 has been shown to bind defective nascent polypeptides, targeting them for aggresome formation ^64^. Its loss may disrupt misfolded protein clearance, compounding aggregation and neurotoxicity. Similar reductions have been reported in the hippocampus of AD patients^65^, reinforcing its relevance to neurodegenerative vulnerability. Our 3xTg model showed stable EEF1A2 expression across timepoints, highlighting possible human specific regulation or context-dependent effects; consistent with findings that SG formation and resolution depend on both intrinsic neuronal properties and environmental stress ^66^.

### 4.4 Transcriptomic Remodeling Reveals Widespread Functional Loss in rpAD

To complement our proteomic findings and further dissect SG composition, we analyzed the associated transcriptome via RNA sequencing. RNA-seq of SG-associated transcripts confirmed enrichment of protein-coding mRNAs and lncRNAs. Notably, rpAD samples exhibited widespread depletion of transcripts encoding RBPs, splicing factors, and cytoskeletal regulators; including DDX17, SFPQ, PABPC1, and EEF1A1. These transcripts are essential for maintaining SG dynamics and mRNA fate ^67,68^. Interestingly, our findings contrast with studies showing increased DDX17 in AD cortex, suggesting possible differential sequestration or cell-type-specific effects ^69^. Physical feature analysis revealed that longer transcripts with extended UTRs and higher GC content were more frequently depleted in rpAD. These features are known to influence translation efficiency and RNA-protein binding, making them more susceptible to decay or misregulation under stress ^70^. The strong overlap between depleted transcripts and known SG-enriched RNAs supports the idea of impaired RNA buffering capacity in rpAD.

### 4.5 Ubiquitin-Proteasome Pathway and SG Interaction in rpAD

To understand the functional implications of these proteomic and transcriptomic shifts, we performed enrichment analyses of the differentially expressed components. Functional enrichment analyses revealed distinct pathway activation between spAD and rpAD. SGs from rpAD brains showed enhanced enrichment for ubiquitin-mediated proteolysis and proteasome components; pathways central to SG turnover and protein quality control ^71^. Dysfunction of the UPS has long been implicated in AD and tau accumulation ^72^, and our findings suggest that SG remodeling may intersect with impaired proteostasis to accelerate neurodegeneration in rpAD. The downregulation of several E3 ligases and proteasome-related enzymes in our rpAD dataset suggests defective protein degradation, potentially contributing to tau and RBP aggregation., This supports a model in which SGs shift from transient stress buffers to chronic reservoirs of misfolded proteins under pathological conditions. Together, these integrated findings provide a unified model of SG remodeling in AD subtypes, as detailed in the conclusion below. This study has limitations, including a small cohort size, reliance on a single cortical region, postmortem variability, and absence of functional validation (e.g., SG manipulation or gene silencing). These findings, while robust, should be interpreted in light of these constraints.

### Conclusion

This study presents the first comprehensive multi-omics profiling of SGs directly isolated from human postmortem brain tissue, revealing profound morphological, proteomic, and transcriptomic remodeling in AD, particularly in its rapidly progressive form (rpAD). Our data show that SGs in rpAD display a striking transformation from small, spherical granules observed in control brains to large, amorphous aggregates, suggestive of impaired disassembly and clearance. Proteomic analysis uncovered a clear shift in SG composition, with control SGs enriched in components related to translation, DNA repair, and ion homeostasis, while SGs from spAD and rpAD were increasingly enriched for proteins involved in MAPK signaling, lipoprotein metabolism, and neuroinflammatory responses. This indicates a pathological reprogramming of SG function during disease progression. Importantly, SGs in rpAD exhibited reduced levels of cytoskeletal and translational regulators, including TUBA1B, TUBA4A, EEF1A2, and HSPA1B, suggesting that compromised structural and translational integrity may underlie SG dysfunction. The presence of tau and its isoforms in SGs, along with altered abundance in rpAD, further implicates SGs in tau aggregation dynamics. At the transcriptome level, rpAD SGs showed a marked depletion of long, GC-rich, protein-coding RNAs;particularly those encoding RBPs and splicing regulators such as DDX17, SFPQ, and PABPC1,indicating reduced RNA buffering capacity. Moreover, the enrichment of pathways related to ubiquitin-mediated proteolysis, RNA splicing, and immune signaling in rpAD highlights the convergence of SG dysfunction with broader disruptions in proteostasis and stress adaptation mechanisms. Together, these findings suggest that SGs are not merely passive responders to cellular stress in AD but might be actively involved in disease acceleration in rpAD, serving as hubs for pathological signaling, tau aggregation, and transcriptomic deregulation. By elucidating disease- and subtype-specific SG alterations, this work opens new avenues for mechanistic understanding and identifies novel candidate targets for therapeutic intervention in aggressive forms of AD

## References

1. Ripin, N. & Parker, R. Formation, function, and pathology of RNP granules. Cell 186, 4737–4756 (2023).

2. Ivanov, P., Kedersha, N. & Anderson, P. Stress granules and processing bodies in translational control. Cold Spring Harb Perspect Biol 11, a032813 (2019).

3. Youn, J.-Y. et al. Properties of stress granule and P-body proteomes. Mol Cell 76, 286– 294 (2019).

4. Adeli, K. Translational control mechanisms in metabolic regulation: critical role of RNA binding proteins, microRNAs, and cytoplasmic RNA granules. American Journal of Physiology-Endocrinology and Metabolism 301, E1051–E1064 (2011).

5. Buchan, J. R. mRNP granules: assembly, function, and connections with disease. RNA Biol 11, 1019–1030 (2014).

6. Riggs, C. L., Kedersha, N., Ivanov, P. & Anderson, P. Mammalian stress granules and P bodies at a glance. J Cell Sci 133, jcs242487 (2020).

7. Vanderweyde, T., Youmans, K., Liu-Yesucevitz, L. & Wolozin, B. Role of stress granules and RNA-binding proteins in neurodegeneration: a mini-review. Gerontology 59, 524– 533 (2013).

8. Kedersha and, N. & Anderson, P. Stress granules: sites of mRNA triage that regulate mRNA stability and translatability. Biochem Soc Trans 30, 963–969 (2002).

9. Protter, D. S. W. & Parker, R. Principles and properties of stress granules. Trends Cell Biol 26, 668–679 (2016).

10. Standart, N. & Weil, D. P-bodies: cytosolic droplets for coordinated mRNA storage. Trends in Genetics 34, 612–626 (2018).

11. Gilks, N. et al. Stress granule assembly is mediated by prion-like aggregation of TIA-1. Mol Biol Cell 15, 5383–5398 (2004).

12. Buchan, J. R. & Parker, R. Eukaryotic stress granules: the ins and outs of translation. Mol Cell 36, 932–941 (2009).

13. Vanderweyde, T., Youmans, K., Liu-Yesucevitz, L. & Wolozin, B. Role of stress granules and RNA-binding proteins in neurodegeneration: a mini-review. Gerontology 59, 524– 533 (2013).

14. Campos-Melo, D., Hawley, Z. C. E., Droppelmann, C. A. & Strong, M. J. The integral role of RNA in stress granule formation and function. Front Cell Dev Biol 9, 621779 (2021).

15. Marcelo, A., Koppenol, R., de Almeida, L. P., Matos, C. A. & Nóbrega, C. Stress granules, RNA-binding proteins and polyglutamine diseases: too much aggregation? Cell Death Dis 12, 592 (2021).

16. Guillén-Boixet, J. et al. RNA-induced conformational switching and clustering of G3BP drive stress granule assembly by condensation. Cell 181, 346–361 (2020).

17. Yang, P. et al. G3BP1 is a tunable switch that triggers phase separation to assemble stress granules. Cell 181, 325–345 (2020).

18. Sidibé, H. & Vande Velde, C. RNA granules and their role in neurodegenerative diseases. The biology of mRNA: Structure and function 195–245 (2019).

19. Mollet, S. et al. Translationally repressed mRNA transiently cycles through stress granules during stress. Mol Biol Cell 19, 4469–4479 (2008).

20. Wolozin, B. & Ivanov, P. Stress granules and neurodegeneration. Nat Rev Neurosci 20, 649–666 (2019).

21. Wang, J., Gan, Y., Cao, J., Dong, X. & Ouyang, W. Pathophysiology of stress granules: An emerging link to diseases. Int J Mol Med 49, 44 (2022).

22. Dudman, J. & Qi, X. Stress granule dysregulation in amyotrophic lateral sclerosis. Front Cell Neurosci 14, 598517 (2020).

23. Fan, A. C. & Leung, A. K. L. RNA granules and diseases: a case study of stress granules in ALS and FTLD. RNA Processing: Disease and Genome-wide Probing 263–296 (2016).

24. Knopman, D. S., et al. Alzheimer disease. Nat Rev Dis Primers 7, 33 (2021).

25. Long, J. M. & Holtzman, D. M. Alzheimer disease: an update on pathobiology and treatment strategies. Cell 179, 312–339 (2019).

26. Hardy, J. & Selkoe, D. J. The amyloid hypothesis of Alzheimer’s disease: progress and problems on the road to therapeutics. Science (1979) 297, 353–356 (2002).

27. Schmidt, C. et al. Rapidly progressive Alzheimer’s disease: a multicenter update. Journal of Alzheimer’s Disease 30, 751–756 (2012).

28. Schmidt, C. et al. Clinical features of rapidly progressive Alzheimer’s disease. Dement Geriatr Cogn Disord 29, 371–378 (2010).

29. Schmidt, C. et al. Rapidly progressive Alzheimer disease. Arch Neurol 68, 1124–1130 (2011).

30. Chitravas, N. et al. Treatable neurological disorders misdiagnosed as Creutzfeldt-Jakob disease. Ann Neurol 70, 437–444 (2011).

31. Schmidt, C. et al. Rapidly progressive Alzheimer disease. Arch Neurol 68, 1124–1130 (2011).

32. Sperling, R. A. et al. Toward defining the preclinical stages of Alzheimer’s disease: Recommendations from the National Institute on Aging-Alzheimer’s Association workgroups on diagnostic guidelines for Alzheimer’s disease. Alzheimer’s & dementia 7, 280–292 (2011).

33. Bisht, K., Sharma, K. & Tremblay, M.-È. Chronic stress as a risk factor for Alzheimer’s disease: roles of microglia-mediated synaptic remodeling, inflammation, and oxidative stress. Neurobiol Stress 9, 9–21 (2018).

34. Liu-Yesucevitz, L. et al. Tar DNA binding protein-43 (TDP-43) associates with stress granules: analysis of cultured cells and pathological brain tissue. PLoS One 5, e13250 (2010).

35. Waelter, S. et al. Accumulation of mutant huntingtin fragments in aggresome-like inclusion bodies as a result of insufficient protein degradation. Mol Biol Cell 12, 1393– 1407 (2001).

36. Johnson, E. C. B. et al. Deep proteomic network analysis of Alzheimer’s disease brain reveals alterations in RNA binding proteins and RNA splicing associated with disease. Mol Neurodegener 13, 1–22 (2018).

37. Vanderweyde, T. et al. Contrasting pathology of the stress granule proteins TIA-1 and G3BP in tauopathies. Journal of Neuroscience 32, 8270–8283 (2012).

38. Zafar, S. et al. Prion protein interactome: identifying novel targets in slowly and rapidly progressive forms of Alzheimer’s disease. Journal of Alzheimer’s Disease 59, 265–275 (2017).

39. Hernández-Ortega, K., Garcia-Esparcia, P., Gil, L., Lucas, J. J. & Ferrer, I. Altered machinery of protein synthesis in Alzheimer’s: from the nucleolus to the ribosome. brain pathology 26, 593–605 (2016).

40. Oddo, S. et al. Triple-transgenic model of Alzheimer’s disease with plaques and tangles: intracellular Aβ and synaptic dysfunction. Neuron 39, 409–421 (2003).

41. Fritzsche, R. et al. Interactome of two diverse RNA granules links mRNA localization to translational repression in neurons. Cell Rep 5, 1749–1762 (2013).

42. Schieweck, R., Ang, F. yee, Fritzsche, R. & Kiebler, M. A. Isolation and characterization of endogenous RNPs from brain tissues. RNA Detection: Methods and Protocols 419– 426 (2018).

43. Nesvizhskii, A. I., Keller, A., Kolker, E. & Aebersold, R. A statistical model for identifying proteins by tandem mass spectrometry. Anal Chem 75, 4646–4658 (2003).

44. Fan, A. C. & Leung, A. K. L. RNA granules and diseases: a case study of stress granules in ALS and FTLD. RNA Processing: Disease and Genome-wide Probing 263–296 (2016).

45. Johnson, E. C. B. et al. Deep proteomic network analysis of Alzheimer’s disease brain reveals alterations in RNA binding proteins and RNA splicing associated with disease. Mol Neurodegener 13, 1–22 (2018).

46. Vanderweyde, T. et al. Contrasting pathology of the stress granule proteins TIA-1 and G3BP in tauopathies. Journal of Neuroscience 32, 8270–8283 (2012).

47. Protter, D. S. W. & Parker, R. Principles and properties of stress granules. Trends Cell Biol 26, 668–679 (2016).

48. Wolozin, B. & Ivanov, P. Stress granules and neurodegeneration. Nat Rev Neurosci 20, 649–666 (2019).

49. Kedersha, N. et al. G3BP–Caprin1–USP10 complexes mediate stress granule condensation and associate with 40S subunits. Journal of Cell Biology 212, (2016).

50. Molliex, A. et al. Phase separation by low complexity domains promotes stress granule assembly and drives pathological fibrillization. Cell 163, 123–133 (2015).

51. Apicco, D. J. et al. Reducing the RNA binding protein TIA1 protects against tau-mediated neurodegeneration in vivo. Nat Neurosci 21, 72–80 (2018).

52. Vanderweyde, T. et al. Interaction of tau with the RNA-binding protein TIA1 regulates tau pathophysiology and toxicity. Cell Rep 15, 1455–1466 (2016).

53. Khong, A. et al. The stress granule transcriptome reveals principles of mRNA accumulation in stress granules. Mol Cell 68, 808–820 (2017).

54. Sato, K., Takayama, K. & Inoue, S. Stress granules sequester Alzheimer’s disease-associated gene transcripts and regulate disease-related neuronal proteostasis. Aging (Albany NY) 15, 3984 (2023).

55. Wolozin, B. & Ivanov, P. Stress granules and neurodegeneration. Nat Rev Neurosci 20, 649–666 (2019).

56. Zhang, P. et al. Chronic optogenetic induction of stress granules is cytotoxic and reveals the evolution of ALS-FTD pathology. Elife 8, e39578 (2019).

57. Molliex, A. et al. Phase separation by low complexity domains promotes stress granule assembly and drives pathological fibrillization. Cell 163, 123–133 (2015).

58. Wolozin, B. Regulated protein aggregation: stress granules and neurodegeneration. Mol Neurodegener 7, 1–12 (2012).

59. Ghosh, S. & Geahlen, R. L. Stress granules modulate SYK to cause microglial cell dysfunction in Alzheimer’s disease. EBioMedicine 2, 1785–1798 (2015).

60. Khong, A. et al. The stress granule transcriptome reveals principles of mRNA accumulation in stress granules. Mol Cell 68, 808–820 (2017).

61. Hu, S. et al. Time-resolved proteomic profiling reveals compositional and functional transitions across the stress granule life cycle. Nat Commun 14, 7782 (2023).

62. Brangwynne, C. P. et al. Germline P granules are liquid droplets that localize by controlled dissolution/condensation. Science (1979) 324, 1729–1732 (2009).

63. Bonifacino, T. et al. Nearly 30 years of animal models to study amyotrophic lateral sclerosis: a historical overview and future perspectives. Int J Mol Sci 22, 12236 (2021).

64. Meriin, A. B., Zaarur, N. & Sherman, M. Y. Association of translation factor eEF1A with defective ribosomal products generates a signal for aggresome formation. J Cell Sci 125, 2665–2674 (2012).

65. Beckelman, B. C. et al. Dysregulation of elongation factor 1A expression is correlated with synaptic plasticity impairments in Alzheimer’s disease. Journal of Alzheimer’s Disease 54, 669–678 (2016).

66. Anderson, P. & Kedersha, N. Stress granules: the Tao of RNA triage. Trends Biochem Sci 33, 141–150 (2008).

67. Khong, A. et al. The stress granule transcriptome reveals principles of mRNA accumulation in stress granules. Mol Cell 68, 808–820 (2017).

68. Tsuiji, H. et al. Spliceosome integrity is defective in the motor neuron diseases ALS and SMA. EMBO Mol Med 5, 221–234 (2013).

69. Liu, Y. et al. DEAD-Box helicase 17 promotes amyloidogenesis by regulating BACE1 translation. Brain Sci 13, 745 (2023).

70. Zheng, D. et al. Predicting the translation efficiency of messenger RNA in mammalian cells. bioRxiv 2024–2028 (2025).

71. Lee, M. J., Lee, J. H. & Rubinsztein, D. C. Tau degradation: The ubiquitin–proteasome system versus the autophagy-lysosome system. Prog Neurobiol 105, 49–59 (2013).

72. Upadhya, S. C. & Hegde, A. N. Role of the ubiquitin proteasome system in Alzheimer’s disease. BMC Biochem 8, 1–8 (2007).

