## Additional Information for "Molecular and structural remodeling of stress granules in slowly and rapidly progressive Alzheimer’s disease"

### Supplementary:

#### Preparation of Human brain homogenate:

Frontal cortex tissues from human brain (cont., spAD, rpAD) were homogenized on ice using brain extraction buffer (EB, 25 mM HEPES [pH 7.3], 150 mM KCl, 8% glycerol, 0.1% NP-40, 1 mM DTT, protease inhibitor cocktail [Roche], 40 U/ml RNase inhibitors [Ribolock, Fermentas]) using homogenizer for 7 minutes. Brain lysates were centrifuged at 20,000x g for 20 minutes at 4C to separate the nuclei and cell debris in the form of pellet and soluble S20 lysate in the form of supernatant. Afterwards, S20 lysate was immediately used for further experiments or frozen at -80C for future experiments.

#### Immunoblotting

To identify the granule marker enriched fraction, we performed western blots for all the samples. Samples of 10µL of each fraction were mixed with 3-µL sample buffer and denatured at 95 °C for 5 minutes. The denatured samples were loaded on SDS gel and run at 100v for 1.5 hours. After the samples were resolved on the gel, it was subjected to Coomassie staining to confirm the successful fractionation. After the confirmation of the fractionation pattern western blot for each sample in the cohort was performed for crucial markers such as TIA-R, PABPC1, Tau-5, T22 and some other markers. Briefly, following electrophoresis and the transfer of proteins on PVDF membrane, it was probed with respective primary antibodies and incubated overnight at 4 °C. Secondary antibodies were probed the next day and developed the membranes to identify the granule-enriched fractions.

##### Gene ontology (GO) term analysis

Gene ontology (GO) term analysis was performed using ShinyGO to identify significantly enriched GO terms related to biological processes, cellular components and molecular functions. The input for analysis consisted of list of proteins identified in the SGs proteome from cont. and disease groups. Background was adjusted with the input of full proteome data identified in the study. Default settings were used unless otherwise specified. GO terms with a false discovery rate (FDR) or p-value below threshold ($p<0.05$) were considered significantly enriched. The results were visualized in the bar plots, highlighting the most significant GO terms.

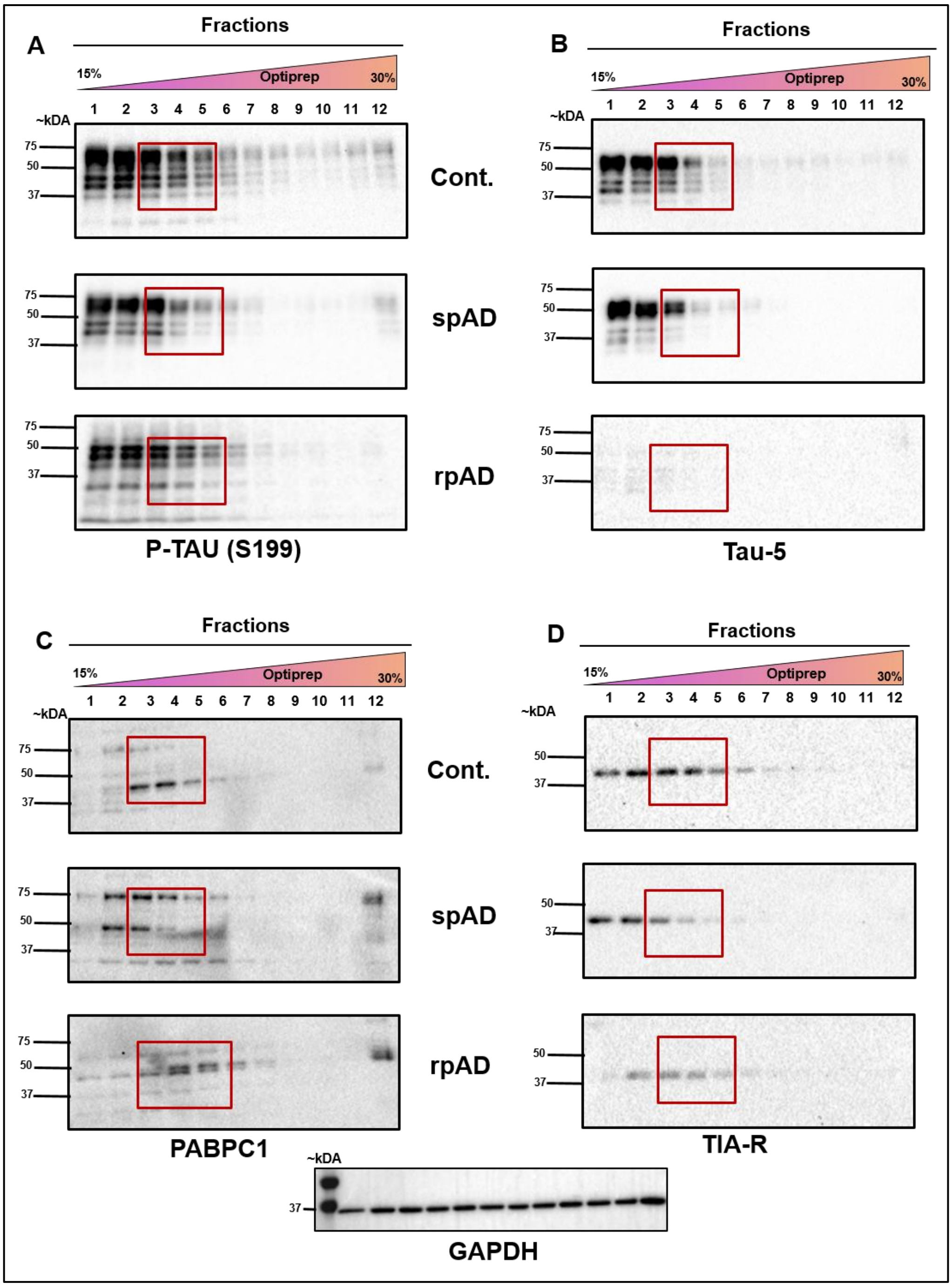


**Figure S1. Western blot analysis of SG markers and tau species in fractionated brain lysates from cont., spAD, and rpAD cases.** (A) Western blot detection of phosphorylated tau (p-Tau S199) across OptiPrep gradient fractions shows disease-dependent enrichment patterns, with strongest signals observed in fractions 3-5. (B) Total tau (Tau-5) distribution reveals similar enrichment in SG-containing fractions (3-5) across all three conditions. (C) Detection PABPC1, a canonical SG marker, confirms co-enrichment in the same gradient fractions, supporting SG association. (D) TIA-R, another well-established SG marker, is also highly enriched in fractions 3–5. Red boxes indicate the fractions with prominent signal intensities for each target protein. These data collectively confirm the localization of both tau species and SG-associated proteins to the same density range of the gradient. n = 15 (Control = 5; spAD = 5; rpAD = 5).

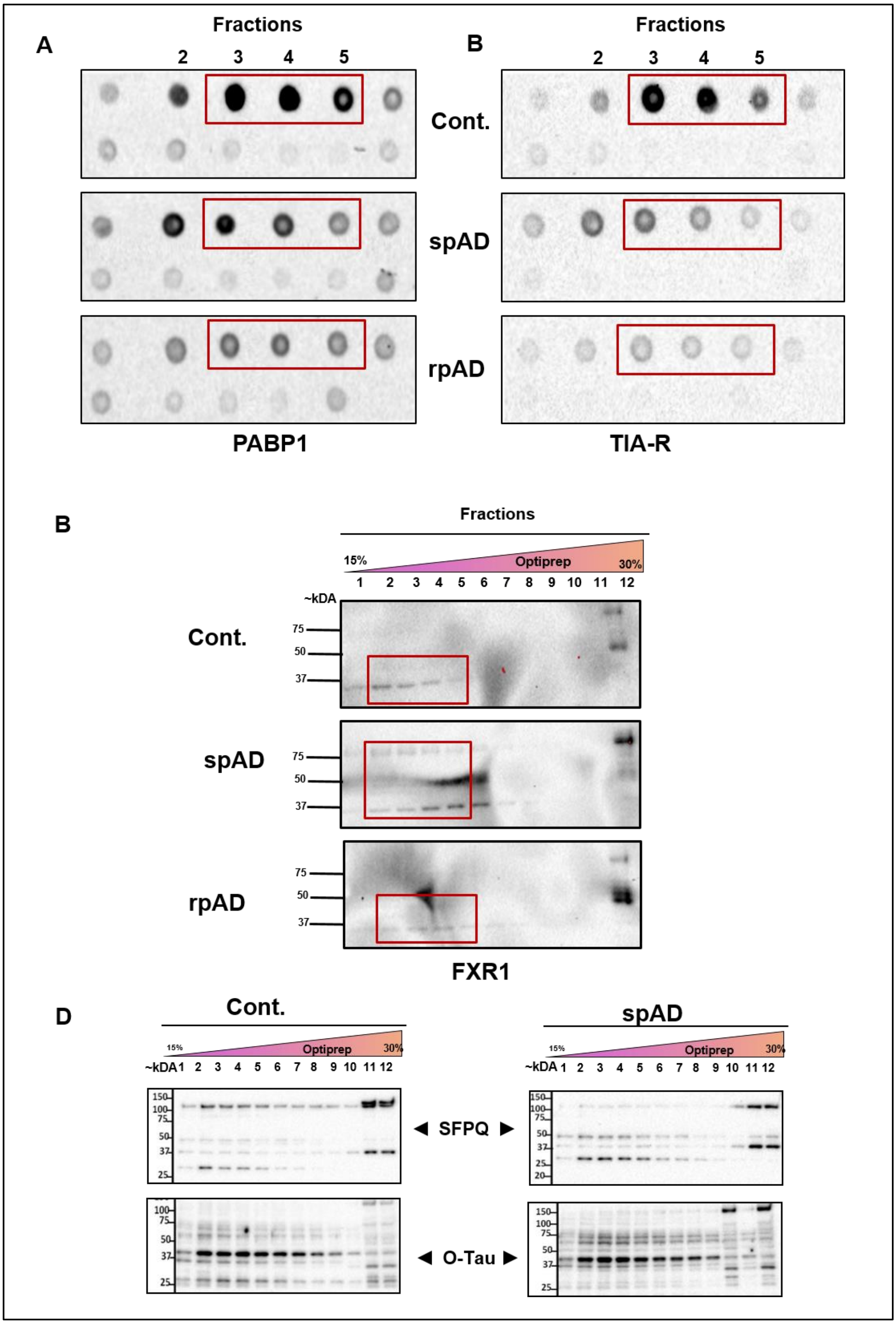


**Figure S2. Dot blot and Western blot analysis of SG markers and AD-related proteins across OptiPrep gradient fractions from control, spAD, and rpAD brain samples**. (A) Dot blot analysis of non-denatured fractions reveals the enrichment of TIA-R across fractions 3–5 in cont., spAD, and rpAD groups, indicating the presence of native SGs. (B) Dot blot analysis of PABPC1 confirms similar enrichment patterns of this canonical SG marker in the same fractions across all conditions. (C) Western blot analysis of denatured fractions shows the distribution of FXR1, SG-RBP, within the SG-enriched fractions (3–5). (D) Western blot analysis reveals co-enrichment of SFPQ and oligomeric tau (O-Tau) in SG-containing fractions from all three groups.

Red boxes highlight the fractions with highest signal intensity. These results validate the association of SG components with specific gradient fractions. n = 15 (Control = 5, spAD = 5, rpAD = 5).

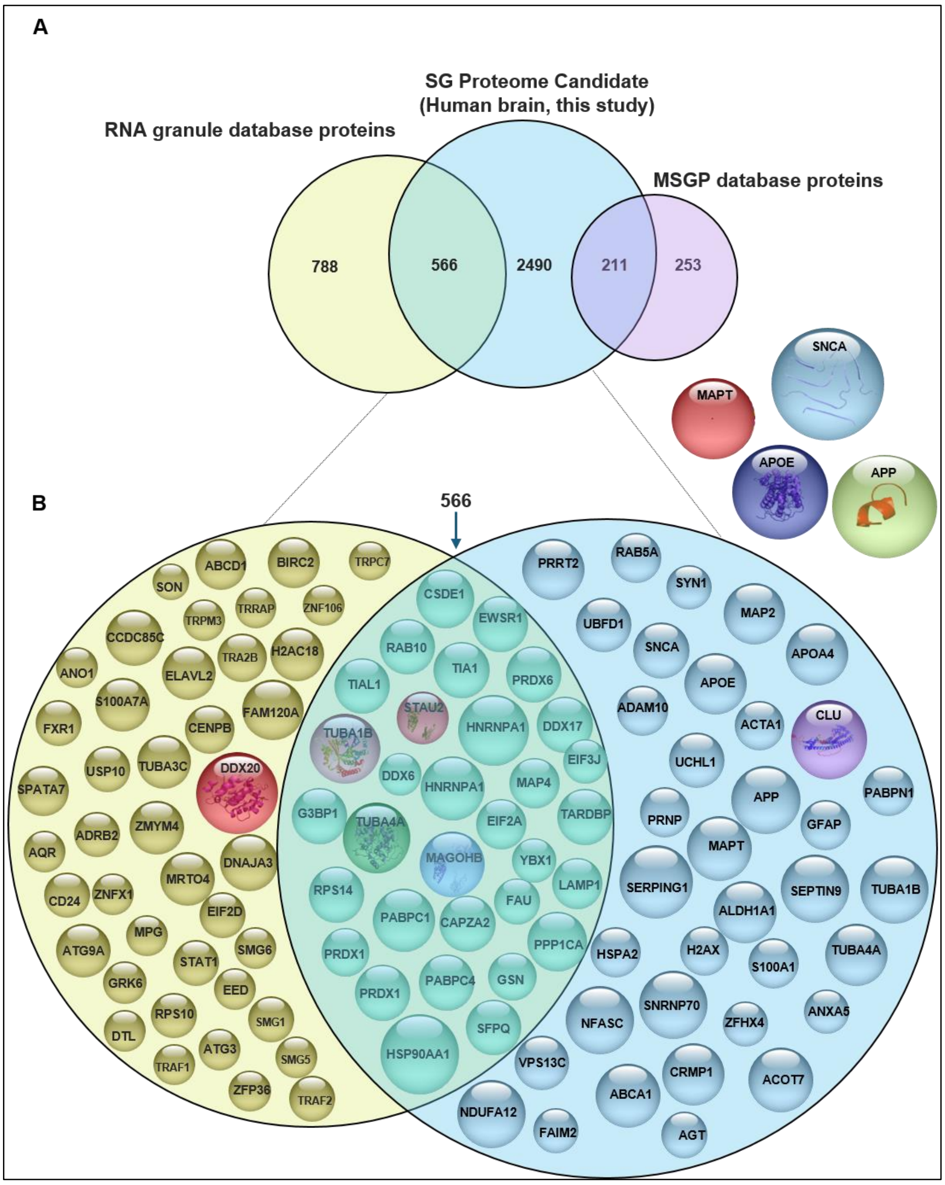


Figure S3: Comparative integration of the human SG proteome with known SG and RNA granule databases. (A) Venn diagram showing the overlap between the SG proteome identified in this study (n = 3,267 proteins) and two reference databases: the RNA granule database (1,354 proteins) and the MSGP database (464 proteins). A total of 566 proteins overlap with the RNA granule database and 211 overlap with the MSGP dataset, highlighting both conserved and uniquely identified SG components from human brain tissue. (B) Bubble plot displaying representative SG-associated proteins detected in this study. Proteins exclusively found in the RNA granule database are shown in yellow, those unique to this study's SG proteome are in blue, and overlapping proteins between both datasets are indicated in green. AD-relevant pathological proteins such as MAPT (tau), SNCA (α-synuclein), APOE, and APP are highlighted outside the main network, emphasizing disease-related SG components. n = 5 per group (cont., spAD, rpAD), each with 3 technical replicates.

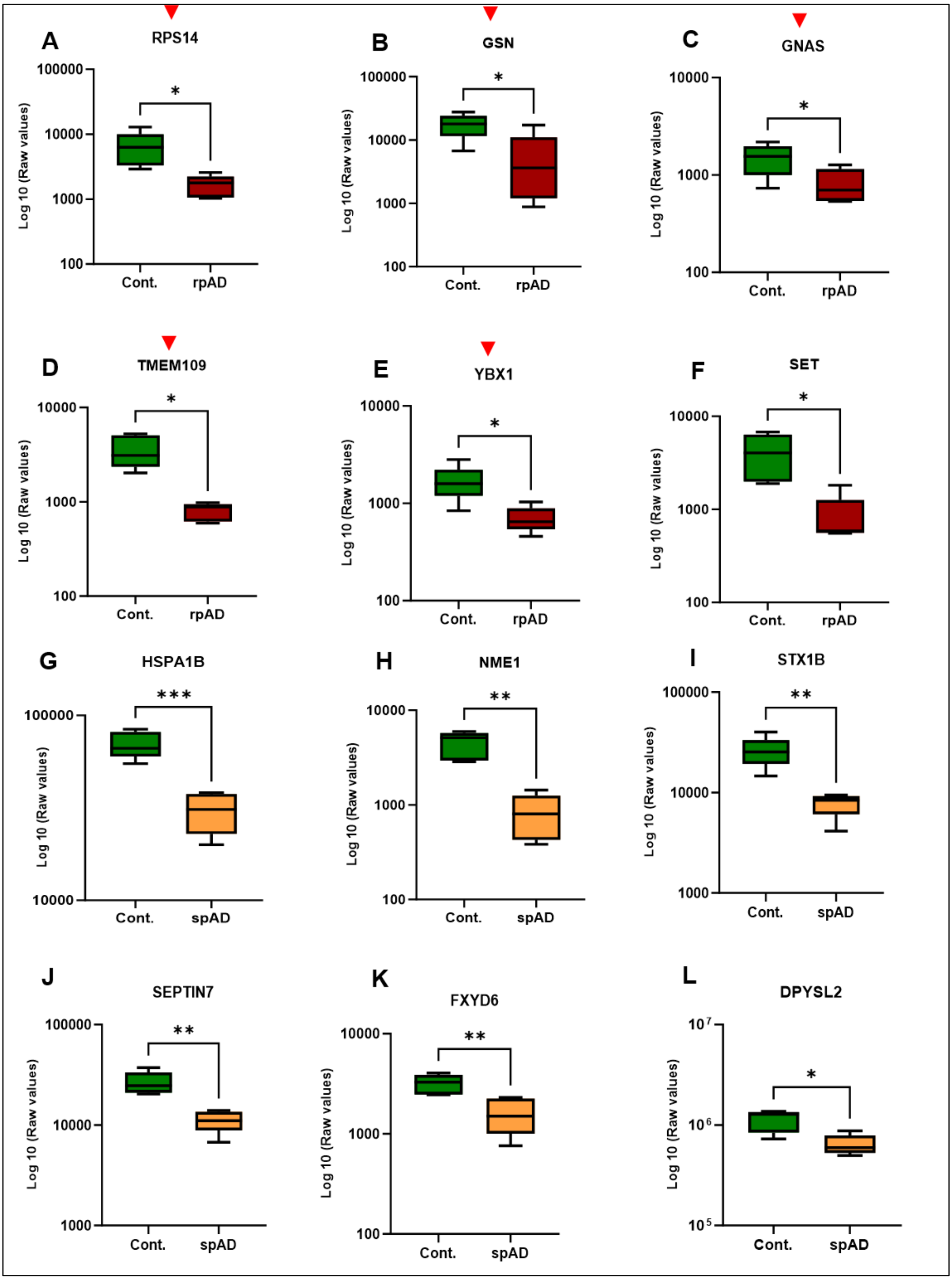


**Figure S4. Quantification of differentially enriched proteins (DEPs) from the SG proteome in Alzheimer's disease subtypes.** (A–F) Box plots showing relative abundance of representative DEPs in SGs between cont. and rpAD groups from MS data. (G–L) Box plots depicting representative DEPs comparing cont. and spAD groups. Each box plot illustrates the statistical distribution of protein abundance based on normalized mass spectrometry data, with RNA granule-associated proteins marked by red arrowheads. Statistical comparisons were performed using Welch's t-test; significance thresholds: $p < 0.05$ (*), $p < 0.01$ (**).

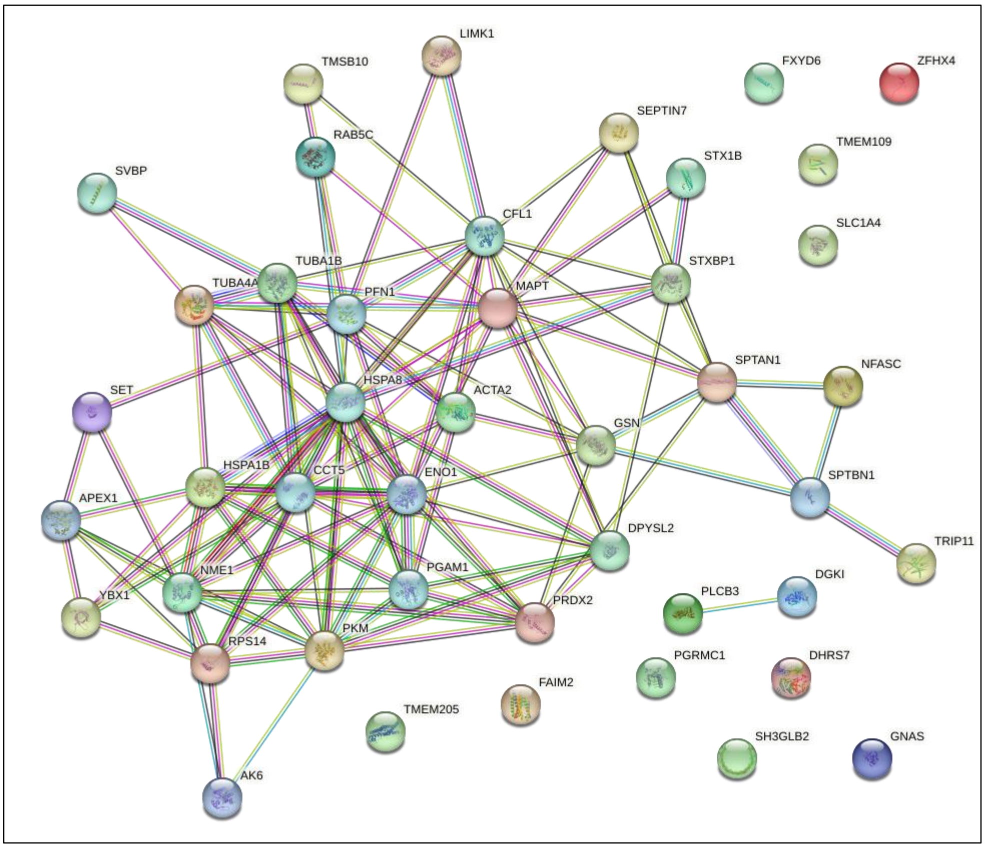


**Figure S5. Protein–protein interaction (PPI) network of DEPs from SG proteome.** A correlation-based PPI network was generated to visualize the functional connectivity among DEPs identified in SGs. Each node represents a protein, while edges denote experimentally validated or predicted functional associations (e.g., physical binding, co-expression, or pathway linkage). Proteins are grouped into interaction clusters, suggesting potential cooperative or pathway-specific functions. Notably, central hubs such as MAPT, TUBA1B, HSPA8, and CCT5 show high connectivity, highlighting their possible regulatory roles within SG-associated molecular networks.

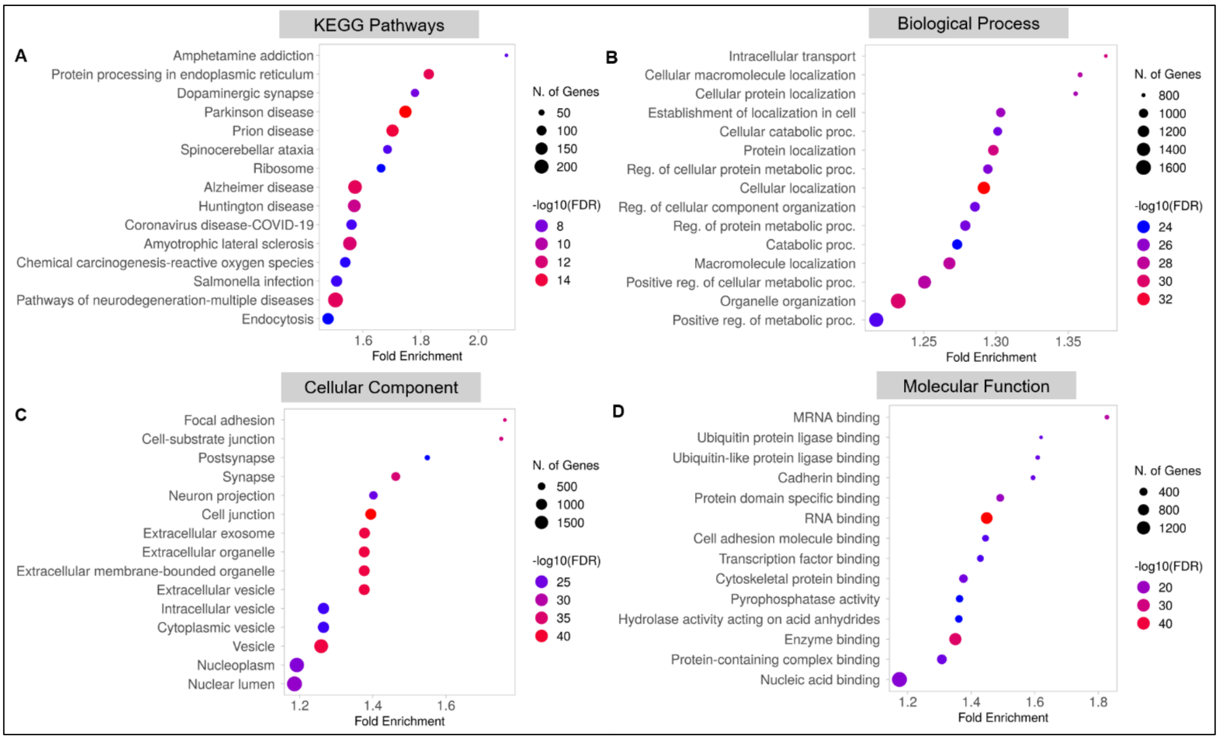


Figure S6: Functional enrichment analysis of SG-associated transcripts. (A) KEGG pathway analysis reveals significant enrichment in pathways related to neurodegenerative diseases, including PD , AD, Huntington's disease, and prion disease, as well as synaptic signaling and endocytosis. (B) GO biological process analysis highlights enriched terms related to intracellular transport, protein localization, cellular component organization, and metabolic regulation, suggesting SG involvement in broad cellular homeostasis mechanisms. (C) GO cellular component analysis identifies predominant localization of SG-associated transcripts in structures such as synapses, neuron projections, cell junctions, exosomes, and various vesicle types. (D) GO molecular function terms are enriched for mRNA binding, RNA binding, ubiquitin ligase activity, and protein complex binding, supporting a role for SGs in regulating RNA metabolism and proteostasis. Dot size represents $-\log_{10}$(FDR) as a measure of statistical significance, while dot color reflects the number of overlapping genes. FDR correction was applied using the Benjamini-Hochberg method via ShinyGO. Fold enrichment quantifies the magnitude of enrichment relative to background transcriptomes.

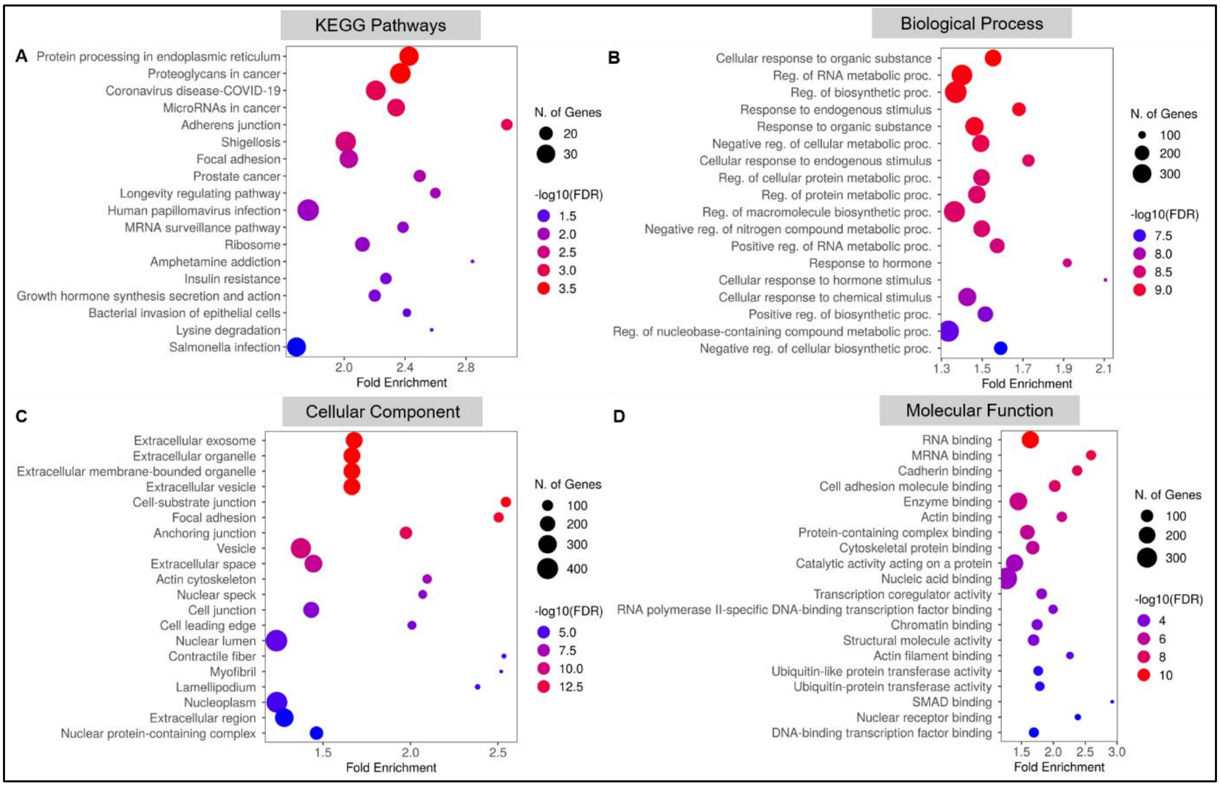


**Figure S7. Functional enrichment analysis of DEGs in the SG-associated transcriptome between cont. and rpAD groups.** (A) KEGG pathway analysis reveals significant enrichment of pathways involved in protein processing, cell adhesion, viral infection, cancer signaling, neurodegeneration and Parkinson's disease. (B) GO biological processes highlight transcriptional and metabolic regulation, responses to chemical and organic stimuli, and RNA metabolic process modulation, suggesting altered cellular stress responses and biosynthetic control in rpAD. (C) GO cellular component analysis shows strong enrichment of DEGs associated with extracellular exosomes, membrane-bounded organelles, vesicles, synapses, and cell junctions- implying altered subcellular localization and structural remodeling. (D) GO molecular function analysis reveals enrichment of RNA binding, cadherin and cell adhesion, molecule binding, protein complex interactions, enzyme binding, and transcription factor activities, indicating disruptions in RNA handling and signaling. Dot size represents the -$\log_{10}$(FDR) as a measure of statistical significance, while dot color denotes the number of overlapping genes per category. Fold enrichment indicates the degree of overrepresentation relative to background. FDR correction was applied using the Benjamini-Hochberg method through ShinyGO.

Table S1. Summary of cases. Control;  spAD: sporadic AD; rpAD: rapid Alzheimer disease; N: normal; Braak NFT: Braak neurofibrillary tangle pathology (0-VI); TAP: Thal αB phase (1-5); CERAD: Consortium to Establish a Registry for Alzheimer disease (C0-C3); NIA score: National Institute on Aging (A0-A3); PMI: post-mortem delay. ABC categorization: A: TAP: amyloid score; B: Baak NFT pathology; C: CERAD (CERAD-NIA-AA score)

| No. | Case | Clinical diagnosis | Age | Gender | Braak NFT | CERAD-(NIAAA score) |
|---|---|---|---|---|---|---|
| 1 | Control 1 | N | 73 | M | - | A0-1B0 |
| 2 | Control 2 | N | 62 | M | - | A0-1 B0 |
| 3 | Control 3 | N | 59 | M | - | A0-1 B0 |
| 4 | Control 4 | N | 62 | M | - | A0-1 B0 |
| 5 | Control 5 | N | 61 | M | - | A0-1 B0 |
| 6 | spAD 1 | AD; amyloid angiopathy | 81 | F | V | C |
| 7 | spAD 2 | AD | 82 | F | V | B |
| 8 | spAD 3 | AD; amyloid angiopathy ;ageing related tau astrogliopathy | 75 | M | V | C |
| 9 | spAD 4 | AD ;LBD amygdala | 82 | M | V | C |
| 10 | spAD 5 | AD, LBD, zerebrale Amyloidangiopathie | 78 | M | V | C |
| 11 | rpAD 1 | AD | 84 | M | V | - |
| 12 | rpAD 2 | AD | 72 | F | VI | - |
| 13 | rpAD 3 | AD, zerebrale Amyloidangiopathie | 71 | F | VI | C |
| 14 | rpAD 4 | AD | 77 | M | VI | - |
| 15 | rpAD 5 | AD | 80 | F | V | C |

**Table S2: List of primary antibodies used in the study**

| Primary Antibody | Origin | Dilution (IB) | Cat. No./ Company |
|---|---|---|---|
| TIA-1 | Rabbit | 1: 500 | ab140595/Abcam |
| TIA-R | Rabbit | 1:1000 | 8509S/ cell siganlling |
| PABPC1 | Rabbit | 1:1000 | 4992S/ cell siganlling |
| Tau-5 | Mouse | 1: 500 | ab80579/Abcam |
| FXR1 | Rabbit | 1:1000 | Abcam |
| P-tau (S199) | Rabbit | 1: 1000 | ab81268/Abcam |
| T22 | Rabbit | 1:1000 | ABN454 |
| SFPQ | Rabbit | 1: 500 | ab38148/Abcam |
| GAPDH | Mouse | 1: 3000 | G8795/Sigma-Aldrich |

**Table S3: Uniquely enriched proteins in SG proteome of spAD**

| ProteinGroups | Genes | Protein Descriptions |
|---|---|---|
| **P00747** | PLG | Plasminogen |
| **P01008** | SERPINC1 | Antithrombin-III |
| **P01042-2** | KNG1 | Isoform LMW of Kininogen-1 |
| **P02652** | APOA2 | Apolipoprotein A-II |
| **P02741** | CRP | C-reactive protein |
| **P02749** | APOH | Beta-2-glycoprotein 1 |
| **P02751** | FN1 | Fibronectin |
| **P02753** | RBP4 | Retinol-binding protein 4;Retinol-binding protein |
| **P02765** | AHSG | Alpha-2-HS-glycoprotein |
| **P04003** | C4BPA | C4b-binding protein alpha chain |
| **P04004** | VTN | Vitronectin |
| **P04040** | CAT | Catalase |
| **P04196** | HRG | Histidine-rich glycoprotein |
| **P04271** | S100B | Protein S100-B |
| **P05546** | SERPIND1 | Heparin cofactor 2 |
| **P06727** | APOA4 | Apolipoprotein A-IV |
| **P06733** | ENO1 | Alpha-enolase |
| **P07360** | C8G | Complement component C8 gamma chain |
| **P07437** | TUBB | Tubulin beta chain |
| **P07741** | APRT | Adenine phosphoribosyltransferase;Isoform 2 of Adenine phosphoribosyltransferase |
| **P08311** | CTSG | Cathepsin G |
| **P09104** | ENO2 | Gamma-enolase |
| **P10768** | ESD | S-formylglutathione hydrolase |
| **P11217** | PYGM | Glycogen phosphorylase, muscle form |
| **P11498** | PC | Pyruvate carboxylase, mitochondrial |
| **P12277** | CKB | Creatine kinase B-type |
| **P14854** | COX6B1 | Cytochrome c oxidase subunit 6B1 |
| **P14868** | DARS1 | Aspartate--tRNA ligase, cytoplasmic |
| **P15259** | PGAM2 | Phosphoglycerate mutase 2 |
| **P15311** | EZR | Ezrin |
| **P16402** | H1-3 | Histone H1.3 |
| **P16403** | H1-2 | Histone H1.2 |
| **P17900** | GM2A | Ganglioside GM2 activator |
| **P18754** | RCC1 | Regulator of chromosome condensation;Isoform 2 of Regulator of chromosome condensation |
| **P21246** | PTN | Pleiotrophin |
| **P22061-2** | PCMT1 | Isoform 2 of Protein-L-isoaspartate(D-aspartate) O-methyltransferase |
| **P23297** | S100A1 | Protein S100-A1 |
| **P24158** | PRTN3 | Myeloblastin |
| **P24588** | AKAP5 | A-kinase anchor protein 5 |
| **P27144** | AK4 | Adenylate kinase 4, mitochondrial |
| **P28907** | CD38 | ADP-ribosyl cyclase/cyclic ADP-ribose hydrolase 1 |
| **P29401** | TKT | Transketolase |
| **P29622** | SERPINA4 | Kallistatin |
| **P29992** | GNA11 | Guanine nucleotide-binding protein subunit alpha-11 |
| **P30048** | PRDX3 | Thioredoxin-dependent peroxide reductase, mitochondrial |
| **P31689** | DNAJA1 | DnaJ homolog subfamily A member 1 |
| **P31949** | S100A11 | Protein S100-A11 |
| **P34897** | SHMT2 | Serine hydroxymethyltransferase, mitochondrial;Isoform 3 of Serine hydroxymethyltransferase, mitochondrial |
| **P34931** | HSPA1L | Heat shock 70 kDa protein 1-like |

| | | |
|---|---|---|
| **P36959** | GMPR | GMP reductase 1 |
| **P38159** | RBMX | RNA-binding motif protein, X chromosome |
| **P40818** | USP8 | Ubiquitin carboxyl-terminal hydrolase 8;Isoform 2 of Ubiquitin carboxyl-terminal hydrolase 8 |
| **P41240** | CSK | Tyrosine-protein kinase CSK |
| **P42025** | ACTR1B | Beta-centractin |
| **P42338** | PIK3CB | Phosphatidylinositol 4,5-bisphosphate 3-kinase catalytic subunit beta isoform |
| **P43686** | PSMC4 | 26S proteasome regulatory subunit 6B;Isoform 2 of 26S proteasome regulatory subunit 6B |
| **P45984** | MAPK9 | Mitogen-activated protein kinase 9;Isoform Alpha-1 of Mitogen-activated protein kinase 9 |
| **P47736-2** | RAP1GAP | Isoform 2 of Rap1 GTPase-activating protein 1 |
| **P47756** | CAPZB | F-actin-capping protein subunit beta |
| **P47756-1** | CAPZB | Isoform 2 of F-actin-capping protein subunit beta |
| **P48454** | PPP3CC | Serine/threonine-protein phosphatase 2B catalytic subunit gamma isoform |
| **P50552** | VASP | Vasodilator-stimulated phosphoprotein |
| **P50570-3** | DNM2 | Isoform 3 of Dynamin-2 |
| **P50991** | CCT4 | T-complex protein 1 subunit delta |
| **P51665** | PSMD7 | 26S proteasome non-ATPase regulatory subunit 7 |
| **P51688** | SGSH | N-sulphoglucosamine sulphohydrolase |
| **P53611** | RABGGTB | Geranylgeranyl transferase type-2 subunit beta |
| **P54652** | HSPA2 | Heat shock-related 70 kDa protein 2 |
| **P55010** | EIF5 | Eukaryotic translation initiation factor 5 |
| **P59768** | GNG2 | Guanine nucleotide-binding protein G(I)/G(S)/G(O) subunit gamma-2 |
| **P62937** | PPIA | Peptidyl-prolyl cis-trans isomerase A |
| **P98179** | RBM3 | RNA-binding protein 3 |
| **Q00535** | CDK5 | Cyclin-dependent kinase 5;Isoform 2 of Cyclin-dependent kinase 5 |
| **Q01968** | OCRL | Inositol polyphosphate 5-phosphatase OCRL;Isoform B of Inositol polyphosphate 5-phosphatase OCRL |
| **Q05193** | DNM1 | Dynamin-1 |
| **Q05329** | GAD2 | Glutamate decarboxylase 2 |
| **Q06203** | PPAT | Amidophosphoribosyltransferase |
| **Q07507** | DPT | Dermatopontin |
| **Q07960** | ARHGAP1 | Rho GTPase-activating protein 1 |
| **Q09028** | RBBP4 | Histone-binding protein RBBP4;Isoform 3 of Histone-binding protein RBBP4 |
| **Q13200** | PSMD2 | 26S proteasome non-ATPase regulatory subunit 2 |
| **Q13409** | DYNC1I2 | Cytoplasmic dynein 1 intermediate chain 2 |
| **Q13424** | SNTA1 | Alpha-1-syntrophin |
| **Q13509** | TUBB3 | Tubulin beta-3 chain |
| **Q14011** | CIRBP | Cold-inducible RNA-binding protein |
| **Q14195** | DPYSL3 | Dihydropyrimidinase-related protein 3 |
| **Q14195-2** | DPYSL3 | Isoform LCRMP-4 of Dihydropyrimidinase-related protein 3 |
| **Q15120** | PDK3 | [Pyruvate dehydrogenase (acetyl-transferring)] kinase isozyme 3, mitochondrial |
| **Q15714** | TSC22D1 | TSC22 domain family protein 1;Isoform 4 of TSC22 domain family protein 1 |
| **Q15811** | ITSN1 | Intersectin-1 |
| **Q16143** | SNCB | Beta-synuclein |
| **Q5EBM0** | CMPK2 | UMP-CMP kinase 2, mitochondrial |
| **Q5RGS4** | PLEKHA1 | Pleckstrin homology domain containing A1 |

| | | |
|---|---|---|
| **Q5T447** | HECTD3 | E3 ubiquitin-protein ligase HECTD3 |
| **Q5TAW7** | CAB39L | Calcium binding protein 39 like (Fragment) |
| **Q5VTT5** | MYOM3 | Myomesin-3 |
| **Q66K74** | MAP1S | Microtubule-associated protein 1S |
| **Q6P1M0** | SLC27A4 | Long-chain fatty acid transport protein 4 |
| **Q6P1Q9** | METTL2B | tRNA N(3)-methylcytidine methyltransferase METTL2B |
| **Q6PUV4** | CPLX2 | Complexin-2 |
| **Q6UXT9** | ABHD15 | Protein ABHD15 |
| **Q7Z6G3** | NECAB2 | N-terminal EF-hand calcium-binding protein 2 |
| **Q7Z6P3** | RAB44 | Ras-related protein Rab-44 |
| **Q86YS7-2** | C2CD5 | Isoform 2 of C2 domain-containing protein 5 |
| **Q8IWA4** | MFN1 | Mitofusin-1 |
| **Q8IYI6** | EXOC8 | Exocyst complex component 8 |
| **Q8IYQ7** | THNSL1 | Threonine synthase-like 1 |
| **Q8N1B4** | VPS52 | Vacuolar protein sorting-associated protein 52 homolog |
| **Q8N2K0** | ABHD12 | Lysophosphatidylserine lipase ABHD12 |
| **Q8N4Q0** | PTGR3 | Prostaglandin reductase 3 |
| **Q8NF37** | LPCAT1 | Lysophosphatidylcholine acyltransferase 1 |
| **Q8TCS8** | PNPT1 | Polyribonucleotide nucleotidyltransferase 1, mitochondrial |
| **Q8WUX9** | CHMP7 | Charged multivesicular body protein 7 |
| **Q92522** | H1-10 | Histone H1.10 |
| **Q92696** | RABGGTA | Geranylgeranyl transferase type-2 subunit alpha |
| **Q92820** | GGH | Gamma-glutamyl hydrolase |
| **Q92930** | RAB8B | Ras-related protein Rab-8B |
| **Q96A00** | PPP1R14A | Protein phosphatase 1 regulatory subunit 14A |
| **Q96A65** | EXOC4 | Exocyst complex component 4 |
| **Q96AB3** | ISOC2 | Isochorismatase domain-containing protein 2 |
| **Q96B36** | AKT1S1 | Proline-rich AKT1 substrate 1 substrate 1 |
| **Q96E17** | RAB3C | Ras-related protein Rab-3C |
| **Q96KP1** | EXOC2 | Exocyst complex component 2 |
| **Q96P70** | IPO9 | Importin-9 |
| **Q96PE3** | INPP4A | Isoform 4 of Inositol polyphosphate-4-phosphatase type I A |
| **Q96RR4** | CAMKK2 | Calcium/calmodulin-dependent protein kinase kinase 2 |
| **Q99584** | S100A13 | Protein S100-A13 |
| **Q99611** | SEPHS2 | Selenide, water dikinase 2 |
| **Q9BPX5** | ARPC5L | Actin-related protein 2/3 complex subunit 5-like protein |
| **Q9BS40** | LXN | Latexin |
| **Q9BUF5** | TUBB6 | Tubulin beta-6 chain |
| **Q9BVA0** | KATNB1 | Katanin p80 WD40 repeat-containing subunit B1 |
| **Q9BW30** | TPPP3 | Tubulin polymerization-promoting protein family member 3 |
| **Q9BXR0** | QTRT1 | Queuine tRNA-ribosyltransferase catalytic subunit 1 |
| **Q9H1E5** | TMX4 | Thioredoxin-related transmembrane protein 4 |
| **Q9H1K0** | RBSN | Rabenosyn-5 |
| **Q9H479** | FN3K | Fructosamine-3-kinase |
| **Q9H492** | MAP1LC3A | Microtubule-associated proteins 1A/1B light chain 3A |
| **Q9H6U6** | BCAS3 | BCAS3 microtubule associated cell migration factor;Isoform 1 of |
| **Q9H9H5** | MAP6D1 | MAP6 domain-containing protein 1 |
| **Q9HB40** | SCPEP1 | Retinoid-inducible serine carboxypeptidase;Isoform 2 of Retinoid-inducible serine carboxypeptidase |
| **Q9NSE4** | IARS2 | Isoleucine--tRNA ligase, mitochondrial |
| **Q9NYQ7** | CELSR3 | Cadherin EGF LAG seven-pass G-type receptor 3 |
| **Q9NZW5** | PALS2 | Protein PALS2 |
| **Q9UHG3** | PCYOX1 | Prenylcysteine oxidase 1 |
| **Q9UID3** | VPS51 | Vacuolar protein sorting-associated protein 51 homolog |
| **Q9UN37** | VPS4A | Vacuolar protein sorting-associated protein 4A |

| | | |
|---|---|---|
| **Q9UPR0** | PLCL2 | Inactive phospholipase C-like protein 2 |
| **Q9UPY6** | WASF3 | Actin-binding protein WASF3 |
| **Q9UQ16-2** | DNM3 | Isoform 2 of Dynamin-3 |
| **Q9Y237** | PIN4 | Peptidyl-prolyl cis-trans isomerase NIMA-interacting 4 |
| **Q9Y265** | RUVBL1 | RuvB-like 1 |
| **Q9Y4E6** | WDR7 | WD repeat-containing protein 7 |
| **Q9Y4Z0** | LSM4 | U6 snRNA-associated Sm-like protein LSm4 |
| **Q9Y512** | SAMM50 | Sorting and assembly machinery component 50 homolog |
| **Q9Y6M5** | SLC30A1 | Proton-coupled zinc antiporter SLC30A1 |

**Table S4: Uniquely enriched proteins in SG proteome of rpAD**

| **Protein Groups** | **Genes** | **Protein Descriptions** |
|---|---|---|
| **P00156** | MT-CYB | Cytochrome b |
| **P00352** | ALDH1A1 | Aldehyde dehydrogenase 1A1 |
| **P00450** | CP | Ceruloplasmin |
| **P00915** | CA1 | Carbonic anhydrase 1 |
| **P00966** | ASS1 | Argininosuccinate synthase |
| **P02042** | HBD | Hemoglobin subunit delta |
| **P02730** | SLC4A1 | Band 3 anion transport protein;Isoform 2 of Band 3 anion transport protein |
| **P02760** | AMBP | Protein AMBP;Alpha-1-microglobulin/bikunin precursor |
| **P02763** | ORM1 | Alpha-1-acid glycoprotein 1 |
| **P03886** | MT-ND1 | NADH-ubiquinone oxidoreductase chain 1 |
| **P05937** | CALB1 | Calbindin |
| **P05976** | MYL1 | Myosin light chain 1/3, skeletal muscle isoform;Isoform MLC3 of Myosin light chain 1/3, skeletal muscle isoform |
| **P06732** | CKM | Creatine kinase M-type |
| **P07196** | NEFL | Neurofilament light polypeptide |
| **P07305** | H1-0 | Histone H1.0 |
| **P08579** | SNRPB2 | U2 small nuclear ribonucleoprotein B |
| **P09012** | SNRPA | U1 small nuclear ribonucleoprotein A |
| **P09936** | UCHL1 | Ubiquitin carboxyl-terminal hydrolase isozyme L1 |
| **P0C870** | JMJD7 | Bifunctional peptidase and (3S)-lysyl hydroxylase JMJD7 |
| **P10586** | PTPRF | Receptor-type tyrosine-protein phosphatase F;Isoform 2 of Receptor-type tyrosine-protein phosphatase F |
| **P10746** | UROS | Uroporphyrinogen-III synthase |
| **P11277** | SPTB | Spectrin beta chain, erythrocytic |
| **P12270** | TPR | Nucleoprotein TPR |
| **P13929** | ENO3 | Beta-enolase |
| **P14625** | HSP90B1 | Endoplasmin |
| **P14866** | HNRNPL | Heterogeneous nuclear ribonucleoprotein L |
| **P15090** | FABP4 | Fatty acid-binding protein, adipocyte |
| **P16219** | ACADS | Short-chain specific acyl-CoA dehydrogenase, mitochondrial |
| **P17302** | GJA1 | Gap junction alpha-1 protein |
| **P18206** | VCL | Vinculin;Isoform 1 of Vinculin |
| **P18669** | PGAM1 | Phosphoglycerate mutase 1 |
| **P19174** | PLCG1 | 1-phosphatidylinositol 4,5-bisphosphate phosphodiesterase gamma-1;Isoform 2 of 1-phosphatidylinositol 4,5-bisphosphate phosphodiesterase gamma-1 |
| **P19429** | TNNI3 | Troponin I, cardiac muscle |
| **P19652** | ORM2 | Alpha-1-acid glycoprotein 2 |
| **P20338** | RAB4A | Ras-related protein Rab-4A |

| | | |
|---|---|---|
| **P21333** | FLNA | Filamin-A;Isoform 2 of Filamin-A |
| **P22314** | UBA1 | Ubiquitin-like modifier-activating enzyme 1;Isoform 2 of Ubiquitin-like modifier-activating enzyme 1 |
| **P24386** | CHM | Rab proteins geranylgeranyltransferase component A 1 |
| **P24534** | EEF1B2 | Elongation factor 1-beta |
| **P25398** | RPS12 | Small ribosomal subunit protein eS12 |
| **P26232** | CTNNA2 | Catenin alpha-2;Isoform 2 of Catenin alpha-2;Isoform 3 of Catenin alpha-2 |
| **P27348** | YWHAQ | 14-3-3 protein theta |
| **P28289** | TMOD1 | Tropomodulin-1;Isoform 2 of Tropomodulin-1 |
| **P28330** | ACADL | Long-chain specific acyl-CoA dehydrogenase, mitochondrial |
| **P30043** | BLVRB | Flavin reductase (NADPH) |
| **P30405** | PPIF | Peptidyl-prolyl cis-trans isomerase F, mitochondrial |
| **P31146** | CORO1A | Coronin-1A |
| **P31153** | MAT2A | S-adenosylmethionine synthase isoform type-2 |
| **P31323** | PRKAR2B | cAMP-dependent protein kinase type II-beta regulatory subunit |
| **P321214** | ARRB2 | Beta-arrestin-2 |
| **P32320** | CDA | Cytidine deaminase |
| **P35520** | CBS | Cystathionine beta-synthase;Isoform 2 of Cystathionine beta-synthase |
| **P42285** | MTREX | Exosome RNA helicase MTR4 |
| **P42765** | ACAA2 | 3-ketoacyl-CoA thiolase, mitochondrial |
| **P43034** | PAFAH1B1 | Platelet-activating factor acetylhydrolase IB subunit beta |
| **P46060** | RANGAP1 | Ran GTPase-activating protein 1 |
| **P46439** | GSTM5 | Glutathione S-transferase Mu 5 |
| **P48147** | PREP | Prolyl endopeptidase |
| **P48723** | HSPA13 | Heat shock 70 kDa protein 13 |
| **P49327** | FASN | Fatty acid synthase |
| **P49588** | AARS1 | Alanine--tRNA ligase, cytoplasmic |
| **P49593** | PPM1F | Protein phosphatase 1F;Isoform 2 of Protein phosphatase 1F |
| **P50213** | IDH3A | Isocitrate dehydrogenase [NAD] subunit alpha, mitochondrial |
| **P50453** | SERPINB9 | Serpin B9 |
| **P50454** | SERPINH1 | Serpin H1 |
| **P50461** | CSRP3 | Cysteine and glycine-rich protein 3 |
| **P50583** | NUDT2 | Bis(5'-nucleosyl)-tetraphosphatase |
| **P52179** | MYOM1 | Myomesin-1 |
| **P52848** | NDST1 | Bifunctional heparan sulfate N-deacetylase/N-sulfotransferase 1 |
| **P53396** | ACLY | ATP-citrate synthase |
| **P53618** | COPB1 | Coatomer subunit beta |
| **P54296** | MYOM2 | Myomesin-2 |
| **P55769** | SNU13 | NHP2-like protein 1 |
| **P55809** | OXCT1 | Succinyl-CoA:3-ketoacid coenzyme A transferase 1, mitochondrial |
| **P61201** | COPS2 | COP9 signalosome complex subunit 2;Isoform 2 of COP9 signalosome complex subunit 2 |
| **P61601** | NCALD | Neurocalcin-delta |
| **P61960** | UFM1 | Ubiquitin-fold modifier 1 |
| **P62072** | TIMM10 | Mitochondrial import inner membrane translocase subunit Tim10 |
| **P62310** | LSM3 | U6 snRNA-associated Sm-like protein LSm3 |
| **P62312** | LSM6 | U6 snRNA-associated Sm-like protein LSm6 |
| **P62714** | PPP2CB | Serine/threonine-protein phosphatase 2A catalytic subunit beta isoform |
| **P62829** | RPL23 | Large ribosomal subunit protein uL14 |

| | | |
|---|---|---|
| **P62877** | RBX1 | E3 ubiquitin-protein ligase RBX1 |
| **P62917** | RPL8 | Large ribosomal subunit protein uL2 |
| **P67936** | TPM4 | Tropomyosin alpha-4 chain |
| **P68871** | HBB | Hemoglobin subunit beta |
| **P69892** | HBG2 | Hemoglobin subunit gamma-2 |
| **P69905** | HBA1 | Hemoglobin subunit alpha |
| **Q03252** | LMNB2 | Lamin-B2 |
| **Q04917** | YWHAH | 14-3-3 protein eta |
| **Q0ZGT2** | NEXN | Nexilin;Isoform 3 of Nexilin;Isoform 4 of Nexilin |
| **Q10567** | AP1B1 | AP-1 complex subunit beta-1 |
| **Q12765** | SCRN1 | Secernin-1 |
| **Q12981** | BNIP1 | Vesicle transport protein SEC20 |
| **Q13033** | STRN3 | Striatin-3;Isoform Alpha of Striatin-3 |
| **Q13045** | FLII | Protein flightless-1 homolog;Isoform 2 of Protein flightless-1 homolog;Isoform 3 of Protein flightless-1 homolog |
| **Q13085** | ACACA | Acetyl-CoA carboxylase 1;Isoform 2 of Acetyl-CoA carboxylase 1 |
| **Q13885** | TUBB2A | Tubulin beta-2A chain |
| **Q14232** | EIF2B1 | Translation initiation factor eIF-2B subunit alpha |
| **Q14249** | ENDOG | Endonuclease G, mitochondrial |
| **Q14289** | PTK2B | Protein-tyrosine kinase 2-beta |
| **Q14315** | FLNC | Filamin-C;Isoform 2 of Filamin-C |
| **Q14642** | INPP5A | Inositol polyphosphate-5-phosphatase A;inositol-polyphosphate 5-phosphatase |
| **Q14692** | BMS1 | Ribosome biogenesis protein BMS1 homolog |
| **Q14696** | MESD | LRP chaperone MESD |
| **Q14697** | GANAB | Neutral alpha-glucosidase AB |
| **Q15018** | ABRAXAS2 | BRISC complex subunit Abraxas 2 |
| **Q15102** | PAFAH1B3 | Platelet-activating factor acetylhydrolase IB subunit alpha1 |
| **Q15119** | PDK2 | [Pyruvate dehydrogenase (acetyl-transferring)] kinase isozyme 2, mitochondrial |
| **Q15124** | PGM5 | Phosphoglucomutase-like protein 5 |
| **Q15149-8** | PLEC | Isoform 8 of Plectin |
| **Q15274** | QPRT | Nicotinate-nucleotide pyrophosphorylase [carboxylating] |
| **Q15459** | SF3A1 | Splicing factor 3A subunit 1 |
| **Q15942** | ZYX | Zyxin |
| **Q16352** | INA | Alpha-internexin |
| **Q16537** | PPP2R5E | Serine/threonine-protein phosphatase 2A 56 kDa regulatory subunit epsilon isoform;Isoform 2 of Serine/threonine-protein phosphatase 2A 56 kDa regulatory subunit epsilon isoform |
| **Q16595** | FXN | Frataxin, mitochondrial |
| **Q16654** | PDK4 | [Pyruvate dehydrogenase (acetyl-transferring)] kinase isozyme 4, mitochondrial |
| **Q3LXA3** | TKFC | Triokinase/FMN cyclase |
| **Q562R1** | ACTBL2 | Beta-actin-like protein 2 |
| **Q5SRE7** | PHYHD1 | Phytanoyl-CoA dioxygenase domain-containing protein 1 |
| **Q5SZR1** | MRPL9 | 39S ribosomal protein L9, mitochondrial;Large ribosomal subunit protein bL9m |
| **Q5T1S4** | RABEPK | Rab9 effector protein with kelch motifs;Rab9 effector protein with kelch motifs (Fragment) |
| **Q5T5C0** | STXBP5 | Syntaxin-binding protein 5 |
| **Q5TDH0** | DDI2 | Protein DDI1 homolog 2;Isoform 3 of Protein DDI1 homolog 2 |
| **Q5U5X0** | LYRM7 | Complex III assembly factor LYRM7 |
| **Q5VVH2** | FKBP1C | Peptidyl-prolyl cis-trans isomerase FKBP1C |
| **Q5VW36** | FOCAD | Focadhesin |

| | | |
|---|---|---|
| **Q5VWZ2** | LYPLAL1 | Lysophospholipase-like protein 1 |
| **Q5W111** | SPRYD7 | SPRY domain-containing protein 7;Isoform 2 of SPRY domain-containing protein 7 |
| **Q69YN2** | CWF19L1 | CWF19-like protein 1;Isoform 3 of CWF19-like protein 1 |
| **Q6ZMU5** | TRIM72 | Tripartite motif-containing protein 72 |
| **Q6ZVK8** | NUDT18 | 8-oxo-dGDP phosphatase NUDT18 |
| **Q702N8** | XIRP1 | Xin actin-binding repeat-containing protein 1 |
| **Q7KZF4** | SND1 | Staphylococcal nuclease domain-containing protein 1 |
| **Q7RTV2** | GSTA5 | Glutathione S-transferase A5 |
| **Q7Z4H8** | POGLUT3 | Protein O-glucosyltransferase 3 |
| **Q7Z7H5** | TMED4 | Transmembrane emp24 domain-containing protein 4 |
| **Q7Z7L7** | ZER1 | Protein zer-1 homolog |
| **Q86UP2** | KTN1 | Kinectin;Isoform 2 of Kinectin;Isoform 4 of Kinectin |
| **Q86WG3** | ATCAY | Caytaxin |
| **Q8IY81** | FTSJ3 | pre-rRNA 2'-O-ribose RNA methyltransferase FTSJ3 |
| **Q8N0X4** | CLYBL | Citramalyl-CoA lyase, mitochondrial;Isoform 2 of Citramalyl-CoA lyase, mitochondrial |
| **Q8N9R8** | SCAI | Protein SCAI;Isoform 2 of Protein SCAI |
| **Q8NFZ4** | NLGN2 | Neuroligin-2 |
| **Q8NHG7** | SVIP | Small VCP/p97-interacting protein |
| **Q8TCJ2** | STT3B | Dolichyl-diphosphooligosaccharide--protein glycosyltransferase subunit STT3B |
| **Q8TDQ7** | GNPDA2 | Glucosamine-6-phosphate isomerase 2 |
| **Q8WU17** | RNF139 | E3 ubiquitin-protein ligase RNF139 |
| **Q8WV74** | NUDT8 | Mitochondrial coenzyme A diphosphatase NUDT8;Isoform 2 of Mitochondrial coenzyme A diphosphatase NUDT8 |
| **Q8WXF1** | PSPC1 | Paraspeckle component 1 |
| **Q8WZ42-6** | TTN | Isoform 6 of Titin |
| **Q92506** | HSD17B8 | (3R)-3-hydroxyacyl-CoA dehydrogenase |
| **Q92523** | CPT1B | Carnitine O-palmitoyltransferase 1, muscle isoform |
| **Q926292** | SGCD | Delta-sarcoglycan;Isoform 2 of Delta-sarcoglycan |
| **Q92673** | SORL1 | Sortilin-related receptor |
| **Q92688** | ANP32B | Acidic leucine-rich nuclear phosphoprotein 32 family member B;Isoform 2 of Acidic leucine-rich nuclear phosphoprotein 32 family member B |
| **Q92736** | RYR2 | Ryanodine receptor 2 Isoform 2 of Ryanodine receptor 2 |
| **Q92743** | HTRA1 | Serine protease HTRA1 |
| **Q92905** | COPS5 | COP9 signalosome complex subunit 5 |
| **Q92973** | TNPO1 | Transportin-1;Isoform 2 of Transportin-1 |
| **Q96EY8** | MMAB | Corrinoid adenosyltransferase MMAB |
| **Q96F81** | DISP1 | Protein dispatched homolog 1 |
| **Q96GK7** | FAHD2A | Fumarylacetoacetate hydrolase domain-containing protein 2A |
| **Q96IX5** | ATP5MK | ATP synthase membrane subunit K, mitochondrial |
| **Q96KR1** | ZFR | Zinc finger RNA-binding protein |
| **Q96PF1** | TGM7 | Protein-glutamine gamma-glutamyltransferase Z |
| **Q96QR8** | PURB | Transcriptional activator protein Pur-beta |
| **Q96RP9** | GFM1 | Elongation factor G, mitochondrial |
| **Q96SU42** | OSBPL9 | Oxysterol-binding protein-related protein 9 |
| **Q99733** | NAP1L4 | Nucleosome assembly protein 1-like 4 |
| **Q9BQ69** | MACROD1 | ADP-ribose glycohydrolase MACROD1 |
| **Q9BT78** | COPS4 | COP9 signalosome complex subunit 4 |
| **Q9BVA1** | TUBB2B | Tubulin beta-2B chain |
| **Q9BWD1** | ACAT2 | Acetyl-CoA acetyltransferase, cytosolic |
| **Q9BY32** | ITPA | Inosine triphosphate pyrophosphatase |
| **Q9BZZ5** | API5 | Apoptosis inhibitor 5 |
| **Q9H0D6** | XRN2 | 5'-3' exoribonuclease 2;Isoform 2 of 5'-3' exoribonuclease 2 |

| | | |
|---|---|---|
| **Q9H1E3** | NUCKS1 | Nuclear ubiquitous casein and cyclin-dependent kinase substrate 1 |
| **Q9H223** | EHD4 | EH domain-containing protein 4 |
| **Q9HAV7** | GRPEL1 | GrpE protein homolog 1, mitochondrial |
| **Q9HBH5** | RDH14 | Retinol dehydrogenase 14 |
| **Q9HCB6** | SPON1 | Spondin-1 |
| **Q9HCJ6** | VAT1L | Synaptic vesicle membrane protein VAT-1 homolog-like |
| **Q9NR19** | ACSS2 | Acetyl-coenzyme A synthetase, cytoplasmic;Isoform 2 of Acetyl-coenzyme A synthetase, cytoplasmic |
| **Q9NRZ7** | AGPAT3 | 1-acyl-sn-glycerol-3-phosphate acyltransferase gamma;Isoform 2 of 1-acyl-sn-glycerol-3-phosphate acyltransferase gamma;Isoform 3 of 1-acyl-sn-glycerol-3-phosphate acyltransferase gamma |
| **Q9NUB1** | ACSS1 | Acetyl-coenzyme A synthetase 2-like, mitochondrial |
| **Q9NVH1** | DNAJC11 | DnaJ homolog subfamily C member 11;Isoform 3 of DnaJ homolog subfamily C member 11 |
| **Q9UJY5** | GGA1 | ADP-ribosylation factor-binding protein GGA1 |
| **Q9ULU8** | CADPS | Calcium-dependent secretion activator 1;Isoform 4 of Calcium-dependent secretion activator 1 |
| **Q9UM19** | HPCAL4 | Hippocalcin-like protein 4 |
| **Q9UMS0** | NFU1 | NFU1 iron-sulfur cluster scaffold homolog, mitochondrial;Isoform 3 of NFU1 iron-sulfur cluster scaffold homolog, mitochondrial |
| **Q9UNS2** | COPS3 | COP9 signalosome complex subunit 3 |
| **Q9Y2Z9** | COQ6 | Ubiquinone biosynthesis monooxygenase COQ6, mitochondrial;Isoform 3 of Ubiquinone biosynthesis monooxygenase COQ6, mitochondrial |
| **Q9Y3I0** | RTCB | RNA-splicing ligase RtcB homolog |
| **Q9Y490** | TLN1 | Talin-1 |
| **Q9Y4G6** | TLN2 | Talin-2 |
| **Q9Y4W6** | AFG3L2 | AFG3-like protein 2 |
| **Q9Y5K8** | ATP6V1D | V-type proton ATPase subunit D |
| **Q9Y5L4** | TIMM13 | Mitochondrial import inner membrane translocase subunit Tim13 |
| **Q9Y5M8** | SRPRB | Signal recognition particle receptor subunit beta |
| **Q9Y646** | CPQ | Carboxypeptidase Q |
